# NOMe-HiC: joint profiling of genetic variants, DNA methylation, chromatin accessibility, and 3D genome in the same DNA molecule

**DOI:** 10.1101/2022.03.29.486102

**Authors:** Hailu Fu, Haizi Zheng, Xiaoting Chen, Matthew T. Weirauch, Louis J. Muglia, Li Wang, Yaping Liu

## Abstract

Cis-regulatory elements coordinate the regulation of their targeted genes’ expression. However, the joint measurement of cis-regulatory elements’ activities and their interactions in spatial proximity is limited by the current sequencing approaches. We describe a method, NOMe-HiC, which simultaneously captures single nucleotide polymorphisms, DNA methylation, chromatin accessibility (GpC methyltransferase footprints), and chromosome conformation changes from the same DNA molecule, together with the transcriptome, in a single assay. NOMe-HiC shows high concordance with state-of-the-art mono-omic assays across different molecular measurements and reveals coordinated chromatin accessibility at distal genomic segments in spatial proximity and novel types of long-range allele-specific chromatin accessibility.

## Background

Cis-regulatory elements (CREs) play instrumental roles in regulating mammalian gene expression[1]. Sets of transcriptional factors (TFs) bring together multiple CREs, which can be up to megabases away, to initiate and maintain the expression of their targeted genes[2]. The development of new methods capable of profiling multiple aspects of gene regulation in the same cell, including gene expression, chromatin conformation, DNA methylation, chromatin accessibility, and genetic variants, remains a longstanding goal and active area of research in the field of functional genomics.

GpC methyltransferase has recently been utilized to footprint TF binding at CREs and reveals the coordinated activities of CREs locally[3]. Since DNA is spatially organized into three-dimensional (3D) space in nuclei, distal CREs can be brought into proximity through chromatin folding[4]. Therefore, it is possible that spatially proximal regions may also exhibit a coordinated GpC methyltransferase footprint. However, conventional GpC methyltransferase based approaches, such as NOMe-seq/dSMF[5,6], have limited power in detecting coordinated CREs activities across large genomic distances due to the short read lengths profiled in next-generation sequencing (NGS). Recently developed SMAC-seq combines methyltransferase with Nanopore sequencing to detect the coordinated activities separated by up to many kilobases[7]. However, this method does not provide the spatial proximity information to distinguish between cis-co-binding events by the same set of TFs and trans-effects.

Moreover, recent studies suggest that allelic differences in genetic content can affect the activity of local CREs by using heterozygous SNPs and their linked CpG methylation or GpC methyltransferase footprint from the same reads[8–10]. In principle, the genetic differences between two alleles may affect the coordinated activities of both CREs that are distant in linear 2D space but proximate spatially in 3D space. However, our current understanding of the long-range allele-specific CRE activities is still limited by both computational approaches and experimental assays. First, most GpC methyltransferase based assays heavily depend on sodium bisulfite treatment, which leads to difficulties in distinguishing C>T SNPs (65% SNPs in dbSNP) and C>T substitutions caused by bisulfite conversion. We previously developed Bis-SNP to identify the SNPs in bisulfite sequencing[11]. However, the genome-wide genotyping accuracy is still not comparable to the whole genome sequencing (WGS). Second, both the short fragment sizes in NGS libraries and the lack of spatial proximity information in long-read sequencing prevent the characterization of spatial interactions between genetic variants and distal CRE activities at the two different alleles.

Chromosome conformation capture (3C) derived technologies, such as *in situ* Hi-C, have been widely used to study 3D genome organization[12]. In Hi-C experiments, spatial proximity information is captured through restriction enzyme digestion and ligation of proximal genomic segments. After the ligation, nucleosomes and TFs are still crosslinked with DNA inside the nuclei, which can be footprinted by the exogenous GpC methyltransferase and detected by the follow-up bisulfite sequencing, similar to the covalent genetic variants and endogenous CpG methylation[13,14]. Therefore, building on our previously established NOMe-seq and Methyl-HiC technologies[5,14], we further developed NOMe-HiC to jointly profile multi-omics from the same DNA molecule, together with the transcriptome in the same assay (Fig. 1).

**Fig.1.**
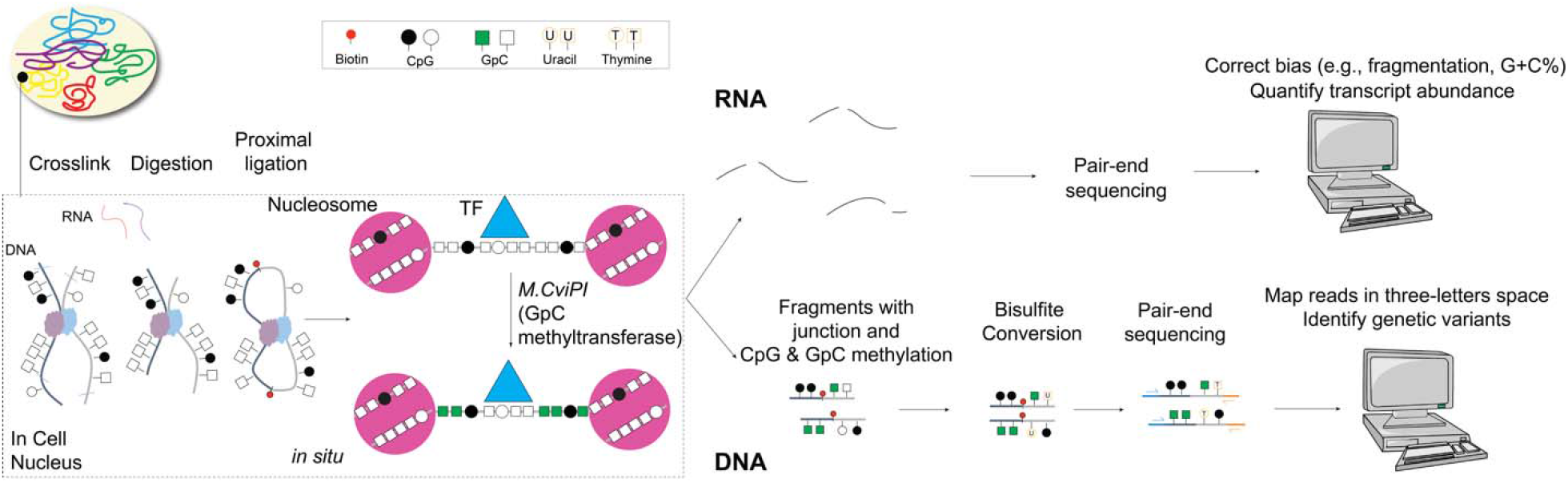
The NOMe-HiC workflow.

## Results

Briefly, after crosslinking, we isolated nuclei and treated them with a methylation insensitive restriction enzyme (e.g., *DpnII*). We use proximity ligation to capture the long-range interactions. We further utilize GpC methyltransferase (*M*.*CviPI*) to footprint chromatin accessibility. After reversing crosslinks, we extract RNA molecules for total RNA-seq. We isolate the DNA molecules with the follow-up bisulfite conversion and paired-end sequencing to obtain covalent endogenous cytosine methylation and exogenous GpC methyltransferase footprint information.

Finally, we improve our previously developed computational method, Bis-SNP, to accurately identify single nucleotide polymorphisms (SNPs) genome-wide from the bisulfite converted reads (Details in Methods, Additional file 1: Supplementary Methods, and Additional file 2: Table S1).

To benchmark the performance, we applied NOMe-HiC to a human lung fibroblast cell line (IMR-90) and a human B-lymphoblastoid cell line (GM12878) (Additional file 2: Table S2-3), which have both been comprehensively profiled publically[15]. First, we compared the measurement of the 3D genome with *in situ* Hi-C results in the same cell lines. The contact matrix is highly similar to that from *in situ* Hi-C (Fig. 2a, Additional file 1: Fig. S1d), with indistinguishable distribution of contact probabilities (Additional file 1: Fig. S1a). Across different resolutions, we also identified comparable compartment score, insulation score, topologically associated domains (TADs), and chromatin loops from the two datasets (Additional file 1: Fig. S1b-c,e-f).

**Fig.2.**
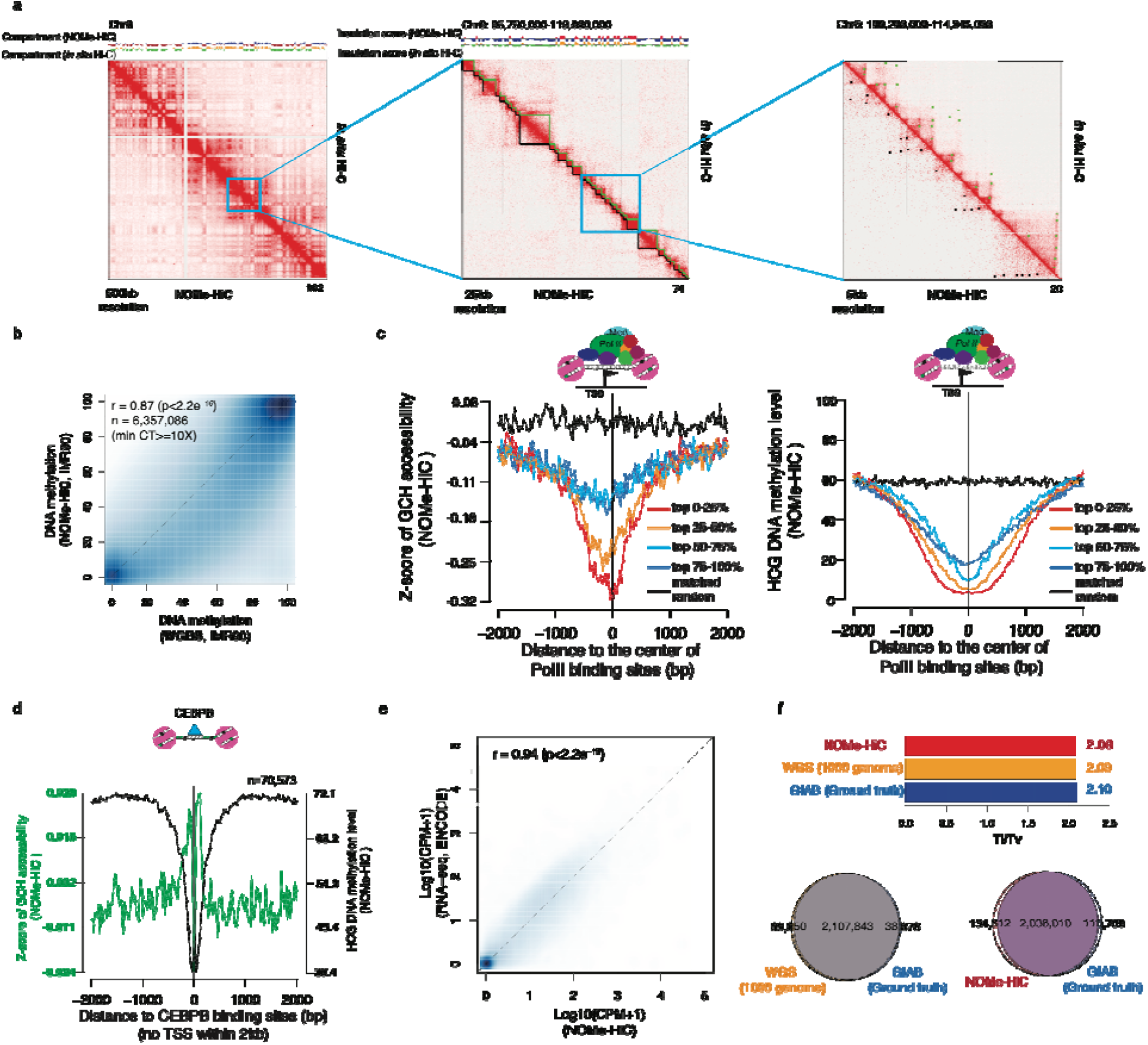
NOMe-HiC generates high-quality multi-omics data. (a). 3D genome concordance with *in situ* Hi-C in IMR-90 cells. From left to right: 500kb resolution for chromosome 6 (including a comparison of the compartment score with Hi-C), 25kb resolution for chromosome 6 (including a comparison of the insulation score and TADs calls with Hi-C), and 5kb resolution for chromosome 6 (including a comparison of chromatin loop calls with Hi-C). (b). Methylation concordance with WGBS at HCG (H=A, C, or T) sites in IMR-90 cells. (c). Average GCH (H=A, C, or T) methyltransferase footprint levels (left) and HCG methylation levels (right) near PolII binding sites. PolII binding sites are divided into quantiles based on the signal strengths of PolII ChIP-seq in IMR-90 cells. Matched random intervals are randomly chosen from the same chromosome and interval lengths as the intervals in the top 0-25% quantile. (d). Average GCH methyltransferase footprint level and HCG methylation level around distal CEBPB binding sites in IMR-90 cells. (e). Scatterplot of concordance of gene expression levels with total RNA-seq levels (ENCODE) in IMR-90 cells. (f). SNPs concordance with the ground truth SNPs data from Genome in a Bottle (GIAB). The results are compared with the SNPs called from WGS (1000 genome project, phase3 in NA12878). Ti/Tv ratio is also compared.

Next, we compared DNA methylation measurements with WGBS from the same cell lines. To avoid the ambiguity of endogenous CpG methylation and exogenous GpC methyltransferase footprint at GpCpG (GCG) sites, we excluded GCG sites for follow-up measurement of CpG methylation and GpC methyltransferase footprint as we did in the NOMe-seq paper[5]. NOMe-HiC captured 31-32 million HCGs (H=A, C, or T) in the human genomes and the methylation levels of these HCGs were highly consistent with those obtained from WGBS (Fig. 2b, Additional file 1: Fig. S1g,i). Similar to Methyl-HiC, NOMe-HiC captured ∼25% fewer HCGs compared to WGBS. Since our previous method NOMe-seq showed residual CpC methylation after GpC methyltransferase treatment, we further excluded CCGs and still found similar high concordance with WGBS in the same cell lines (Additional file 1: Fig. S1h).

Next, we tested the measurement of GpC methyltransferase footprints by using GCH accessibility levels (H=A, C, or T) at TFs binding sites. Notably, nucleosomes and TFs are both crosslinked with DNA. Part of the open chromatin regions will still show a low GCH accessibility level when TFs are bound, which is different from conventional NOMe-seq without the fixations. We observed a concordant lower level of GCH methyltransferase footprints at PolII binding sites with a higher TF occupancy level measured by PolII ChIP-seq in the same cell line (Fig. 2c, Additional file 1: Fig. S2). We also observed the expected GCH footprint patterns and HCG methylation level at TF binding sites that usually bind distantly to the transcriptio starting sites (TSS) (Fig. 2d, Additional file 1: Fig. S3). Further, we utilized the same PCR primers in the previous NOMe-seq study[5] to validate the local accessibility level at individual known opened (CTCF peak), closed (LAMB3), and imprinting regions (SNRPN) and obtained the expected accessibility and DNA methylation results, as reported previously (Additional file 1: Fig. S4). Notably, the accessibility pattern differences between the two cell lines within the CTCF peak showed concordant differences of signal strength obtained from CTCF ChIP-seq (Additional file 1: Fig. S4b).

We next compared the measurement of the transcriptome with total RNA-seq from the same cell lines. Messenger RNA (mRNA) is highly fragmented after the formaldehyde fixation step (crosslinking). After computational correction of the possible biases resulting from fragmentation and G+C%, the transcriptome we obtained is generally concordant with that obtained from total mRNA-seq, in terms of absolute quantification (Fig. 2e, Additional file 1: Fig. S1j) and differentially expressed genes (Additional file 1: Fig. S5).

Finally, we compared the measurement of SNPs (GM12878) with deep WGS from the 1000 genome project performed on the same individual (NA12878, a.k.a. HG001). We evaluated the performance of each using the ground truth genotype results obtained from the same individual by the Genome in a Bottle Consortium (GIAB), which are well characterized through the integration of multiple sequencing technologies. Results with improved Bis-SNP at NOMe-HiC showed similar high performance to that obtained from WGS (Fig.2f, Additional file 1: Fig. S1k) and very high accuracy at sites with callable genotypes (Additional file 1: Fig. S1l). Taken together, these results demonstrate that NOMe-HiC can simultaneously and accurately profile the 3D genome, DNA methylation, GCH methyltransferase footprints (chromatin accessibility), the transcriptome, and SNPs in biological samples (Additional file 1: Fig. S6).

To ensure the robustness of the NOMe-HiC technology, we performed multiple steps of experimental and computational quality controls (QC) (Details in Methods, Additional file 1: Supplementary Methods, and Additional file 1: Fig. S7). We measured the reproducibility of the method by comparing the similarity of different molecule measurements across biological replicates from the same cell lines and found consistently high reproducible results across all of the molecule measurements in both cell lines (Additional file 1: Fig. S8).

We next asked if the genomic regions linearly separated but positioned proximately in 3D space would have coordinated epigenetic statuses. To this end, we analyzed the GCH methyltransferase footprint obtained from NOMe-HiC reads in separated chromatin loop anchors (Fig. 3a). Indeed, the GCH methyltransferase footprint in NOMe-HiC read pairs from the same DNA molecules mapping to separated loop anchors showed a similarly high correlation to that obtained from local regions (Fig. 3b). To ensure that the concordance obtained from the same DNA molecules is not simply due to similar levels of GCH methyltransferase footprints between the loop anchors, we shuffled the links of read pairs while maintaining the average GCH methyltransferase footprint at the same loop anchors. The shuffled read pairs indeed showed significantly less correlation (Fig. 3a-c, Fisher’s (1925) z test, p<2.2e^-16^). We repeated the same analysis across biological replicates at the two different cell types. The GCH methyltransferase footprint showed consistently high long-range epigenetic coordination effects at spatial proximal regions across replicates but with some variation between cell types, suggesting potential cell type specificity (Fig. 3b, Additional file 1: Fig. S9). In addition to the exogenous GCH methyltransferase footprint, we also calculated the spatial coordination of endogenous DNA methylation levels in the two cell lines by using the methylation levels obtained from WCGs (W=A or T). The results showed a significantly higher correlation obtained from the same DNA molecules in spatial proximal regions to that obtained from different DNA molecules in the same regions, which is consistent with our previous Methyl-HiC study on mouse embryonic stem cells (mESCs)[14] (Additional file 1: Fig. S10). However, different from our previous study in mESCs, the long-range coordination of DNA methylation here is significantly lower than the local correlation obtained from these two human cell lines, suggesting a possible variation of long-range endogenous DNA methylation coordination across species and/or cell types. In the human reference genome, GCHs (∼1 in 13bp) measured by NOMe-HiC showed a much higher density than WCGs (∼1 in 125bp) and HCGs (∼1 in 83bp), which provides a new opportunity to study the epigenetic coordination of CREs (∼10-100bp scale) in spatial proximal regions (Additional file 1: Fig. S11). However, the coverage in our most highly sequenced cell line (IMR-90) is still limited (∼20kb resolution in 3D genome). Future studies with much deeper sequenced libraries (and thus with higher resolution), together with long-read sequencing technologies, will likely provide a comprehensive overview of CRE coordination over large genomic distances.

**Fig.3.**
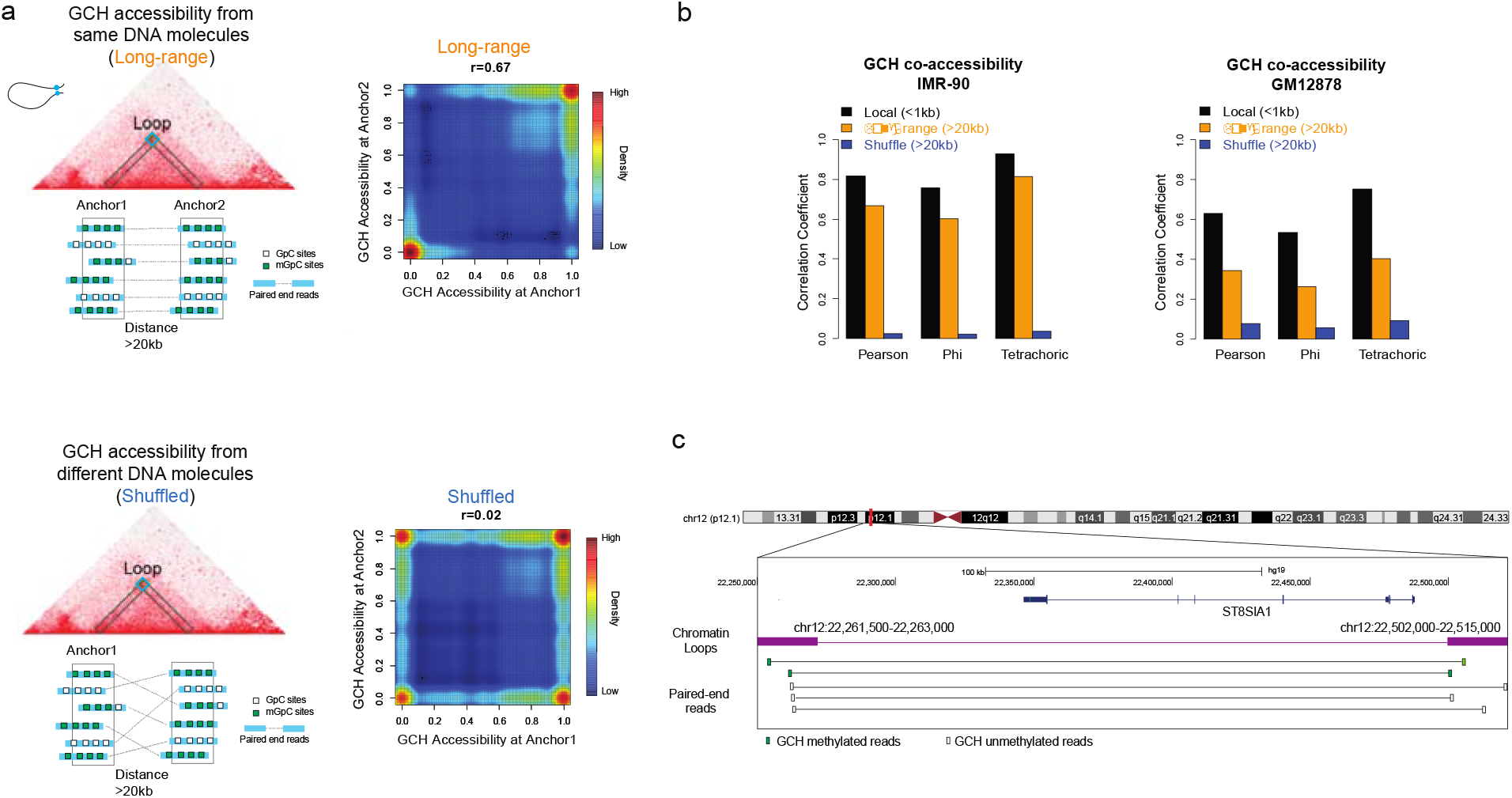
NOMe-HiC reveals coordinated GCH methyltransferase footprints at distal regions in spatial proximity. (a). The coordinated GCH methyltransferase footprint is much stronger at read pairs from the same DNA molecules (top panel) than read pairs from different DNA molecules within the same regions (bottom panel). Each dot in the heatmap represents one read pair within the chromatin loop anchor regions. (b). GCH methyltransferase concordance levels, measured by Pearson Correlation Coefficient, Phi Correlation Coefficient, and Tetrachoric correlation, across all chromatin loop anchor regions (>20kb distance between anchors, orange color, n=125,293 in IMR-90 and n=28,464 in GM12878), their matched local regions (utilizing the read pairs from the same anchor and within 1kb distance, black color, n=378,687 in IMR-90 and n=701,019 in GM12878), and matched shuffled read pairs from the same chromatin loop anchor regions (>20kb distance between anchors, blue color).

We further utilized NOMe-HiC to analyze the long-range allele-specific GCH methyltransferase footprint. We separated the distantly connected NOMe-HiC read pairs into two groups by their parent-of-origin at the heterozygous SNPs overlapped in one end (SNP anchors, Fig. 4a). Since the NOMe-HiC read pairs mostly come from the same DNA molecules, the other ends of the read pairs were also separated into two alleles (although they do not necessarily overlap with heterozygous SNPs) and mapped over large genomic distances to the SNP anchors (non-SNP anchors, Fig. 4a). In this manner, we are able to analyze the long-range allele-specific GCH methyltransferase footprint. We identified three groups of allele-specific footprint loci: those with GCH accessibility differences between the two alleles at both anchors (Group 1, n=1,901, FDR<0.05 at both anchors, Additional file 2: Table S4), only at SNP anchors (Group 2, n=2,941, FDR<0.05, Additional file 2: Table S4), or only at non-SNP anchors (Group 3, n=2,627, FDR<0.05) (Fig. 4a, Additional file 1: Fig. S12, Additional file 2: Table S4). To validate if the allele-specific GCH methyltransferase footprint indeed reflects the allelic imbalance of the CRE activities, we utilized allele-specific TF bindings events identified by applying the MARIO method[16] to TF ChIP-seq reads in GM12878 cells (Details in Methods). We found that allele-specific TF binding sites from ChIP-seq reads are significantly enriched in the SNP anchors of allele-specific GCH methyltransferase footprint sites from NOMe-HiC compared to random permutation controls (Fisher’s exact test, p<2.2e^-16^, Additional file 1: Fig. S13). Further, to characterize the function of these long-rang allele-specific footprints, we checked the enrichment of the anchors over known TF binding sites. Group 1 showed the highest enrichment for CTCF binding sites at both anchors (Fig. 4b, upper panel), suggesting that disruption of the CTCF binding by genetic variants at the SNP anchor may also affect the loop structure, and thus, the binding activities at the non-SNP anchor in spatial proximity. Group 2 exhibited the highest enrichment of CTCF and PolII binding sites at the SNP anchors, but enrichment of different TFs at the non-SNP anchors, which usually bind to distal enhancer regions (Fig. 4b, middle panel). Group 3 showed the highest depletion of TF binding sites that are usually bound to the promoter regions (Fig. 4b, bottom panel).

**Fig.4.**
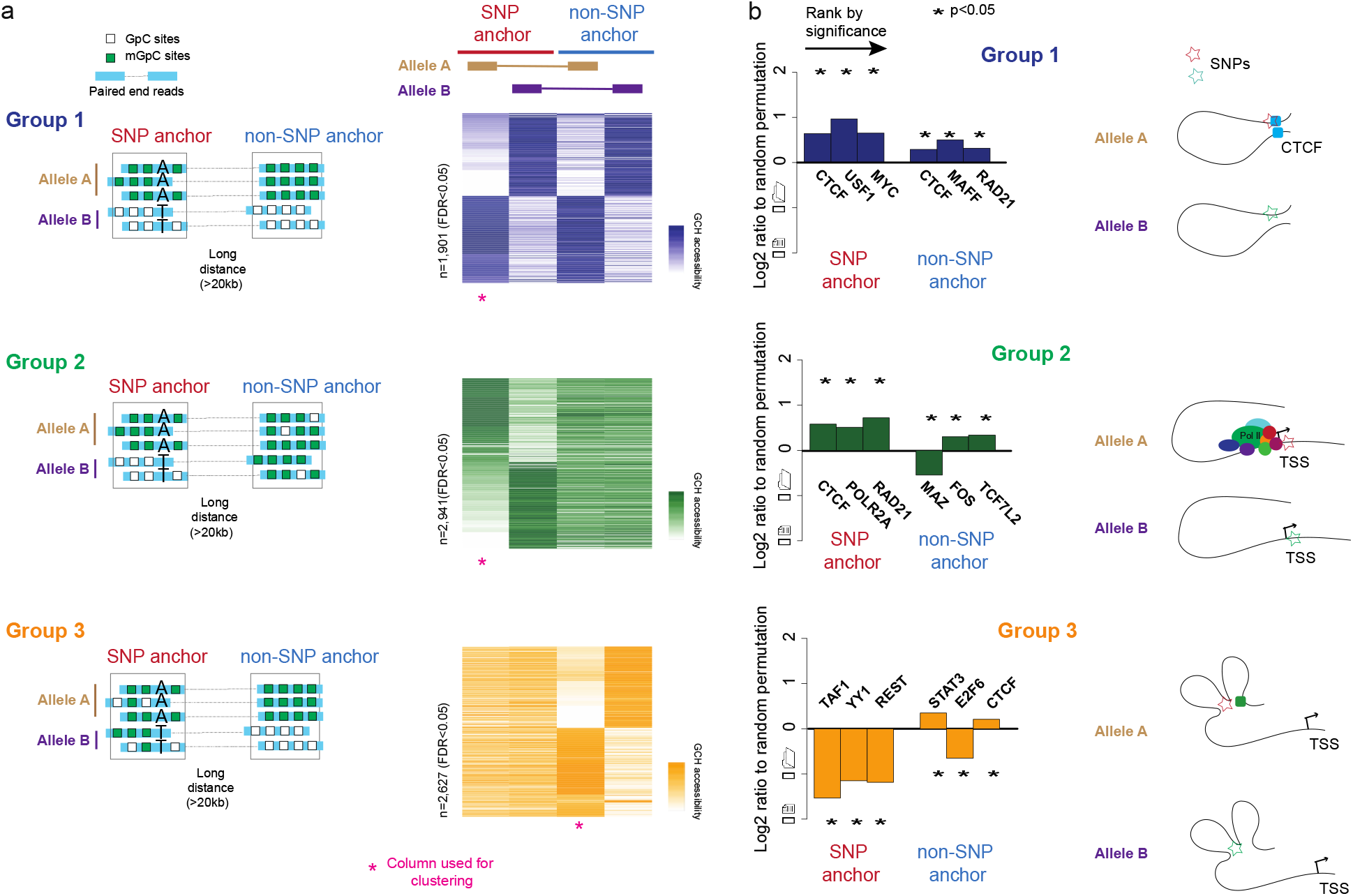
NOMe-HiC reveals the long-range allele-specific GCH methyltransferase footprint. (a). Scheme and heatmap representations of the three groups’ long-range GCH methyltransferase footprint. The distance between SNP anchors and non-SNP anchors is required to be more than 20kb. Group 1 (purple) is the group of loci with significant differences of GCH methyltransferase footprint between Allele A and Allele B at both SNP anchors (FDR<0.05) and non-SNP anchors (FDR<0.05). Group 2 (green) is the group of loci with significant differences in GCH methyltransferase footprint between Allele A and Allele B only at SNP anchors (FDR<0.05) but not at non-SNP anchors (FDR>0.95). Group 3 (orange) is the group of loci with significant differences of GCH methyltransferase footprints between Allele A and Allele B only at non-SNP anchors (FDR<0.05) but not at SNP anchors (FDR>0.95). The rows of the heatmap are clustered by columns marked with the magenta star (*). (b). The log2 enrichment of TF binding sites at SNP anchors and non-SNP anchors. The log2 ratio enrichment level is calculated by the ratio to the matched random intervals in the same chromosomes. The random permutation is repeated ten times. Only the top three enriched TFs are shown.

Genome-wide association studies (GWAS) have associated common complex diseases with hundreds of thousands of genetic variants. Due to linkage disequilibrium (LD), many SNPs near the tag SNPs in a GWAS study are also identified to be significantly associated with the traits. The local CRE activities, especially the imbalanced CREs activities linked to the switching of risk and non-risk alleles nearby, are often utilized to prioritize potential causal SNPs within LD blocks. The Group 3 long-range allele-specific footprint suggests that the switching of risk and non-risk alleles does not need to affect the local chromatin accessibility but instead the activities of distant CREs in spatial proximity. Indeed, we observed the overlap of SNP anchors at Group 3 in GM12878 (B-lymphoblastoid cell) with four LD blocks that are significantly associated with inflammatory bowel disease (IBD)[17,18](Figure 5, complete list is shown in Additional file 2: Table S5). Here, instead of methylation or accessibility QTLs mapping, which require multiple samples, NOMe-HiC provides an alternative approach to link SNPs with the allelic imbalance of distal CREs activities in spatial proximity in a single assay. Future targeted base studies are needed to increase the coverage at regions of interest for further validation,

**Fig.5.**
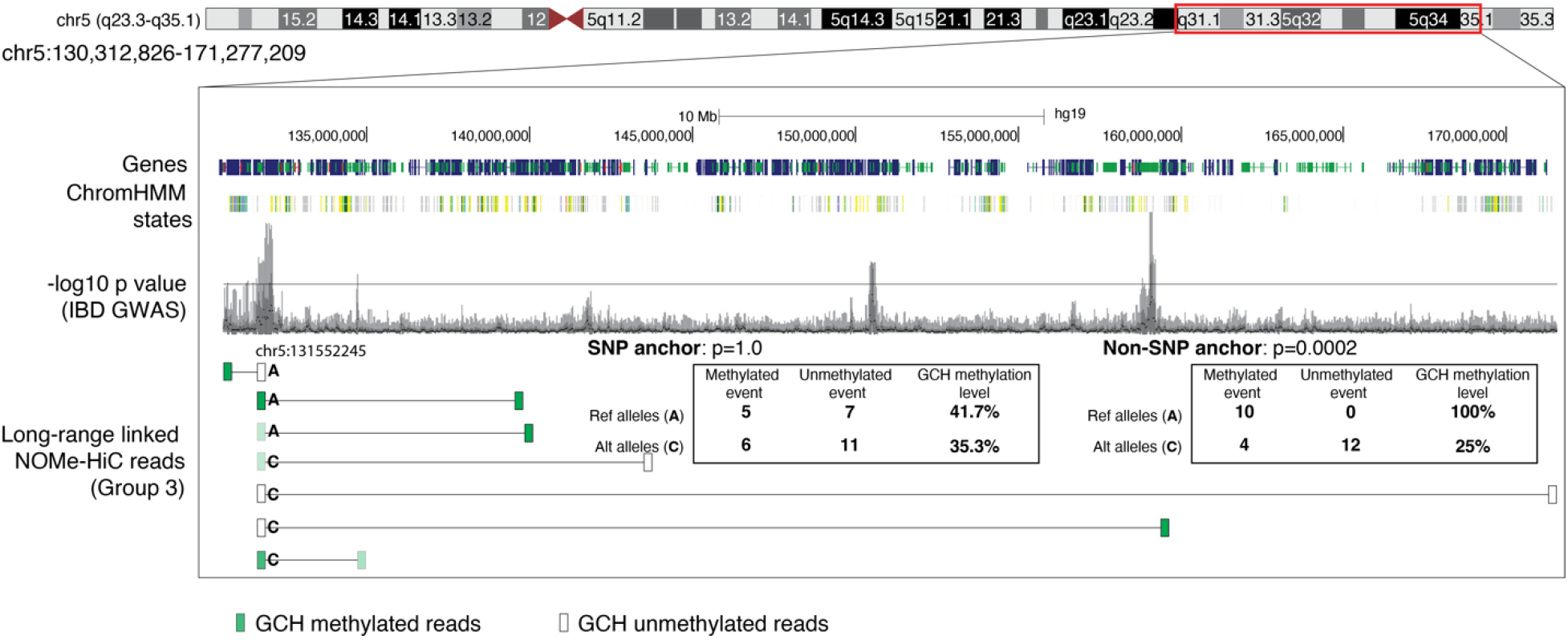
Example of how to use the long-range allele-specific GCH methyltransferase footprint to annotate the allelic imbalance activities of GWAS variants. ChromHMM states in GM12878 were downloaded from the NIH Epigenome Roadmap. -log10 p value of IBD GWAS results are obtained from https://www.ibdgenetics.org/. Only high quality long-range linked reads (>20kb distance) that overlapped with the chr5:13155245 locus are shown in the figure. The transparency of green color represents the methylation level of GCH in the reads: 100% Opacity represent 100% GCH methylation in the reads. Fisher’s Exact test was utilized to calculate the p value of GCH methylation differences between reference alleles and alternative alleles in SNP anchors and non-SNP anchors.

## Discussion

Our NOMe-HiC approach here allows us to characterize the epigenetic status over large genomic distances but are spatially closed. We observed that spatial proximal regions are epigenetic concordant with each other, in terms of both DNA methylation and chromatin accessibility/GpC methyltransferase footprint. The differences in genetic variations between two alleles may not indicate the local allelic imbalanced epigenetic status but, instead, the long-range allelic activities that are in spatial proximity.

There are some limitations to our current genome-wide approach to understanding the concordance between specific pairs of CREs. First, even after merging several sequencing libraries, the resolution is still low (∼20kb resolution) to understand the dynamics at individual paired CREs (10-100bp scale). Second, this is a sequencing approach to pooled cells. For paired regions over large genomic distances, only DNA ligated (in spatial proximity) are profiled. Therefore, the dynamic interactions between paired CREs over large genomic distances are not fully captured across the cell populations. For example, in paired TF1 and TF2 over large genomic distances, only TF1 and TF2 that is spatially close in the sub-group of cells are mostly profiled. Due to the cellular heterogeneity, for the same paired regions (TF1-TF2) that do not have interactions in the other sub-groups of cells, we cannot get the linked epigenetic status between them. Long-read sequencing that covered these two regions in the same DNA molecules (sampling from all cells) combined with our NOMe-HiC technology (sampling mostly from cells that have interactions between two distant regions) may provide an overview of how often these two CREs are concordant with each other across cells. NOMe-seq/dSMF utilized the paired-end reads sampling from all of the cells in the specific local regions, which can be utilized to estimate the relative concordance level between two paired CREs, which, however, have to be linearly close to each other due to the limitation of short reads. Moreover, the library complexity in the current NOMe-HiC protocol is still limited. This limitation may be overcome in the future by bisulfite-free library construction strategies[19,20].

We developed a computational approach to improve the accuracy of our previously developed Bis-SNP method for the detection of genetic variants in bisulfite-converted reads, which can be applied to not only Methyl-HiC or NOMe-HiC but also other bisulfite sequencing data, such as WGBS, and provided a comparable genome-wide accuracy as WGS.

The future extension of NOMe-HiC together with the long-read sequencing technology to the single-cell level will eventually allow us to further characterize long-range gene-regulatory mechanisms from genetic variants to molecular phenotypes, within heterogeneous tissues.

## Conclusions

Overall, we have developed a method to simultaneously study the genetic variants, DNA methylation, GCH methyltransferase footprints (chromatin accessibility), and the 3D genome in the same DNA molecule, together with the transcriptome, all in the same assay. Combining multiple assays allowed us to uncover the coordinated chromatin accessibility/GCH methyltransferase footprints and long-range allele-specific footprints at genomic regions that are far away in linear DNA sequences but spatially close in the nucleus.

## Methods

### Cell culture

The human lung fibroblast cell line IMR-90 was purchased from ATCC and cultured in EMEM (ATCC) with 10% FBS (Gibco) and 1% penicillin/streptomycin. After reaching 70∼80 confluence, cells were detached with 0.5% Trypsin and harvested by centrifuge at 300g for 5min. The human B-lymphoblastoid cell line GM12878 was obtained from CORIELL INSTITUTE and cultured with RPMI-1640 supplemented with 15% FBS and 1% penicillin/streptomycin. GM12878 cells were harvested at a density around 0.7M to 0.8M/mL by centrifuge at 300g for 5min.

### NOMe-HiC

Cells were crosslinked with 1% formaldehyde for 10 min at room temperature, then quenched by adding 0.2M Glycine. After washing with ice-cold PBS, cells were suspended in the nuclei isolation buffer with RNase inhibitors and incubated on ice for 1hr. Subsequent to nuclei membrane permineralization, chromatin was digested using 100U *DpnII* (NEB, R0543L) overnight at 37 □ with RNase inhibitors. The next day, DpnII was inactivated, and nuclei suspension was cooled to room temperature and subjected to biotin fill-in. Proximal segments were *in situ* ligated using T4 DNA ligase (NEB, M0202) supplemented with RNase inhibitors. Then GpC methyltransferase footprint was performed by incubating the nuclei with *M*.*CviPI* (NEB, M0227L), and PBS was added to stop the reaction. After that, one-third of the nuclei were separated for RNA isolation using MagMAX™ FFPE DNA/RNA Ultra Kit (Thermo A31881). The total RNA library was prepared using SMARTer® Stranded Total RNA-Seq Kit v2 (TAKARA 634413). The rest of the nuclei were reverse crosslinked, DNA was precipitated and sonicated to average 400bp followed by 1xAMPure beads cleanup. Biotin pulldown was performed using streptavidin beads, DNA on beads was end-repaired, d-A tailed, and ligated with truncated cytosine-methylated X-gene Universal Stubby Adapter (IDT). 0.1%∼0.5% unmethylated lambda DNA was spiked-in. Bisulfite conversion was conducted using the EZ DNA Methylation-Gold Kit (D5006). Bisulfite converted DNA library was amplified using truncated TruSeq™–Compatible Indexing Primer and KAPA HiFi Uracil+ ReadyMix (Roche, KK2801), supplemented with MgCl_2_. Library quality was determined by qPCR and BioAnalyzer 2000 (Agilent Technologies). Pooling of multiplexed sequencing samples, clustering, and sequencing was carried out as recommended by the manufacturer on Illumina HiSeq XTen or NovaSeq S6000 with the 150 paired-end. RNA-seq libraries or Phi-X (20%) were spiked in to overcome the imbalance of GC ratio of the bisulfite-converted DNA libraries.

### NOMe-HiC (DNA) data analysis

Raw reads were first trimmed using Trim Galore! (v0.6.6, with Cutadapt v2.10)[21] with parameters “--paired_end --clip_R1 5 --clip_R2 5 --three_prime_clip_R1 5 -- three_prime_clip_R2 5” to remove adapters and low-quality reads. The clipped length was determined based on the composition of base pairs along the sequencing cycle by FastQC (v0.11.9). The reference genome (b37, human_g1k_v37.fa) was *in silico* converted to make a C/T reference (all Cs were converted to Ts) and G/A reference (all Gs were converted to As). Paired-end reads were mapped by our previously developed bhmem (v0.37)[14] to each converted reference. Only uniquely mapped and mapping quality passed reads (mapQ>30) on both ends were joined. Reads marked as PCR duplicated reads were filtered. The reads with incompletely converted cytosine (WCH, W=A or T, H=A, C, or T) were filtered out as previously described[11]. Joint read pairs with more than 20kb insertion size were considered as long-range interactions and subjected to the following interaction analysis. Bam files were further converted to .hic files for the following 3D genome analysis similar to Hi-C data (sam2juicer.py) (details in the analysis of *in situ* Hi-C data). HCG methylation and GCH accessibility level were calculated by bissnp_easy_usage.pl with Bis-SNP (v0.90) in NOMe-seq mode. Only bases with a quality score of more than 5 were included in the downstream methylation analysis. More details were implemented in Bhmem.java with parameters “-outputMateDiffChr -buffer 100000”.

### NOMe-HiC (RNA) data analysis

Raw reads were first trimmed as paired-end reads using Trim Galore! (v0.6.6, with Cutadapt v2.10) with parameters “--paired_end --clip_R1 15 --clip_R2 15 --three_prime_clip_R1 5 -- three_prime_clip_R2 5” to remove the adapters and low-quality reads. The clipped length was determined based on the composition of base pairs along the sequencing cycle by FastQC (v0.11.9). Both alignment-based (STAR, v2.7.6a[22], and RNA-SeQC, v2.3.5[23]) and alignment-free (Salmon, v1.5.1[24]) approaches were tried. Salmon was finally used for the bias correction, visualization, and data analysis due to the slightly better concordance with RNA-seq data from ENCODE. The reference transcriptome was obtained from Gencode (v33lift37) and indexed by Salmon with k=31. The parameters used for Salmon quantification were: “quant -- seqBias --gcBias --posBias -l A --validateMappings”. Differential expression analysis was performed on a gene-level by the edgeR (3.32.1)[25] package in R (4.0.5). Only genes with FDR<0.05 and log2 fold change > 1 were considered as the differentially expressed genes between IMR-90 and GM12878. The same RNA-seq analysis pipeline was applied to the total RNA-seq data from ENCODE.

### Quality Controls (QC) for NOMe-HiC

To control for under- or over-treatment, we examined GCH accessibility at several negative and positive control regions (Additional file 1: Fig. S4). We followed the same treatment time and enzyme concentration as that in the original NOMe-seq paper - the accessibility at control regions was largely the same as before. To make sure the nuclei are still in good shape, we examined nuclei status after each step (see Additional file 1: Figure s7): cross-linking, restriction enzyme digestion, biotin fill-in, ligation, and GpC methyltransferase treatment. After bisulfite conversion, we measured DNA quantity and quality with BioAnalyzer 2000 before and after library construction. Before deep sequencing, we sequenced the library to shallow coverage using the Illumina MiSeq platform to check the quality of the DNA and RNA libraries. In computational analyses, we examined the QC metrics for each molecule measurement, as was done in the previous studies: (1). For DNA methylation, we examined the bisulfite conversion rate at non-CpG site in autosomes, mitochondria DNA, and spike-in lambda DNA. (2). For GCH accessibility, we examined GCH methyltransferase efficiency and endogenous DNA methylation levels at important regulatory elements, such as CGI promoters and CTCF binding sites. We also examined the average GCH accessibility globally in autosome and mitochondria DNA. (3). For 3D genome measurements, we examined the fraction of cis-long-range interactions and trans-interactions. (4). For genetic variant calling, we examined the Ti/Tv ratio. (5). For RNA, we utilized RNASeQC2[26] to summarize the RNA QC. (6). We also examined the unique mapping efficiency, PCR duplication rate, and library complexity, as all other common NGS data metrics using PICARD, FastQC, and MultiQC[27] tools.

### Analysis of WGBS data

DNA methylation levels in WGBS were directly downloaded from ENCODE (see details in Additional file 2: Table S1) and extracted from the sites that overlapped with HCGs or WCGs in the reference genome.

### Analysis of *in situ* Hi-C data

Loops were identified by MUSTACHE[28] with parameters “-r 25000 -norm KR -d 5000000”. TopDom[29] was used to call TAD domains, with parameter “window.size = 5” and input KR normalized matrix from strawr[30] (resolution 50k). FAN-C (v0.9.21)[31] was used to calculate insulation scores, with subcommand insulation and parameters “@10kb@KR -w 500000”. To calculate A/B compartment scores, we first used the eigenvector command in Juicer tools[32] (v1.22.01) to obtain the first eigenvector of the Hi-C observed/expected correlation matrix, with parameters KR and BP 500000. Then we calculated the average GC contents for each 500kb bin. With the GC contents and the eigenvector aligned, we flipped the sign of the eigenvector if needed so that high GC content regions are associated with positive compartment scores (compartment A), and low GC content regions are associated with negative compartment scores (compartment B).

### Accurate identification of SNPs from NOMe-HiC

BisulfiteGenotyper in Bis-SNP (v0.90)[11] was utilized to identify SNPs with a genotype quality score of more than 20 at NOMe-HiC data (raw genotype). Further, VCFpostprocess was utilized to filter out poor quality SNPs (regions with sequencing depth over 250X, regions with strand bias more than -0.02, regions with sequencing depth more than 40X and fraction of reads with mapping quality 0 more than 0.1, regions with genotype quality divided by sequencing depth less than 1, and regions with 2 SNPs nearby within +/-10bp). Finally, the resulting SNPs were utilized for imputation and phasing by Minimac4[33] (v1.0.2) and Eagle[34] (v2.4.1) together with the 1000G phase3 v5 panel (EUR population). Since NA12878 was already profiled in the 1000G phase3 project, to avoid potential biases, NA12878 was excluded from the panel for the imputation and phasing. Only biallelic SNPs loci were utilized for imputation and phasing. Other parameters during the SNP imputation and phasing followed similar steps to those recommended by the Michigan Imputation Server[35]. After imputation, only biallelic SNPs with minor allele frequency more than 1% and R^2 more than 0.3 were kept for downstream analysis. WGS genotyping results (NA12878) were downloaded from the 1000G project (phase3)[36].

Gold standard genotype results (NA12878, a.k.a HG001) were downloaded from the Genome in a Bottle (GIAB)[37] website. Bcftools (v1.10.2)[38] was utilized to summarize the Ti/Tv ratio. *Concordance* in GATK (v4.1.9.0)[39] was utilized to evaluate concordance among gold standard (GIAB), WGS (1000G), and NOMe-HiC.

### Correlation analysis of GCH methyltransferase footprints in NOMe-HiC

Only read pairs with at least three GCH sites at both ends were kept for analysis. For the regions of interest, such as chromatin loop anchor regions, Pearson Correlation Coefficient (PCC) was calculated directly from the GCH methylation level of each read of the read pair, not by the average GCH methylation level at each end of anchor regions. Only read pairs spanning at least a 20kb genomic distance were considered in the analysis. As a control, all of the read pairs within the same genomic regions, no matter whether they have long-range interaction or not, were randomly shuffled 100 times and then subjected to PCC calculation. Fisher’s (1925) z, implemented in “cocor.indep.groups” at R package “cocor”[40], was used to assess the significance between the observed PCC from interacting read pairs and PCC of the random read pairs. The scripts for long-range analysis are provided in MethyCorAcrossHiccups.java with parameters “-methyPatternSearch GCH -methyPatternCytPos 2 -bsConvPatternSearch WCH -bsConvPatternCytPos 2 -useBadMate -useGeneralBedPe”. For the short-range mode, to reduce the time cost, we randomly sampled the read pairs up to 100 pairs in each local region with the additional parameters “-onlyShort 1000 -randomSampleReads 100”. Since the methylation levels are largely binary, we also tried to binarize the data (GCH methylation<0.5 is 0 and GCH methylation>=0.5 is 1) and applied the Phi Correlation Coefficient and Tetrachoric correlation by using the psych package (v 2.2.9) in R (v4.2.1).

### Analysis of long-range allele-specific GCH methyltransferase footprints in NOMe-HiC

Only read pairs with one end overlapping with heterozygous SNPs, and GCH methylation levels at both ends were kept for the analysis. At each heterozygous SNP position with enough reads covered at both the reference and alternative alleles, a p-value was calculated by Fisher’s Exact test and followed by FDR correction (Benjamini-Hochberg). Only loci with FDR<0.05 were considered as significant allele-specific loci. For the SNP anchor-Only and non-SNP anchor-Only groups, the methylation differences between alleles should have FDR>0.95 at the insignificant anchor. The scripts for this analysis are provided in LongRangeAsm.java with parameters “-minDist 20000 -coverageRef 2 -coverageAlt 2 -methyPatternSearch GCH - methyPatternCytPos 2 -methyPattern2ndSearch HCG -methyPattern2ndCytPos 2 - bsConvPatternSearch WCH -bsConvPatternCytPos 2 -useBadMate -minMapQ 30”. LD blocks were calculated at 1000 Genome Phase3 v5 autosome VCF files by BigLD with r^2>0.8 [41].

### Validation of local chromatin accessibility and DNA methylation at individual regions

To validate that NOMe-HiC shows a similar GCH methyltransferase footprint at individual regions to that observed using NOMe-seq, we amplified the targeted regions with the same primer sequences from the previous NOMe-seq study [5]. Briefly, DNA materials were collected after the bisulfite conversion step of the NOMe-HiC workflow. For each PCR reaction, 16ng bisulfite converted DNA was obtained and amplified via KAPA HiFi Uracil+ Kit (Roche 7959052001) in a total volume of 25uL, with 300nM primers under the following conditions: 95°C 3min; 35 to 45 cycles of 98°C for 20s, 56°C for 30s, and 72°C for 60s; followed by 72°C for 5min. A 0.8X AMPure bead purification was performed. The PCR products’ sizes ranged from 150 to 600bp by BioAnalyzer (Agilent). 0.715 ng of purified DNA from each region was pooled together, resulting in a total of 5 ng. Library construction was performed on the 5 ng of pooled PCR amplicons using the KAPA HyperPrep Kit (Roche 07962363001) and NEXTFLEX Unique Dual Index Barcodes (7.5µM final concentration, PerkinElmer 514153). Libraries were sequenced on an Illumina MiSeq in PE150 mode.

### Allele-specific TF binding in ChIP-seq data from GM12878 cells

To identify allelic imbalance of sequencing reads overlapping heterozygous genetic variants, we applied the MARIO pipeline v3.93[16] to a collection of 573 ChIP-seq datasets performed in the GM12878 EBV-immortalized B cell line. In brief, MARIO identifies common genetic variants that are (1) heterozygous in the assayed cell line and (2) located within a peak in a given ChIP-seq dataset. MARIO examines the sequencing reads that map to each heterozygote in each peak for imbalance between the two alleles. Within the MARIO pipeline, all ChIP-seq datasets were first processed by removing duplicate reads using the “MarkDuplicates” tool from the PICARD package (https://broadinstitute.github.io/picard/) and then mapped to a masked hg19 genome using STAR aligner [22]. The masked genome was created by changing all common genetic variants to N in the standard hg19 genome, which removes potential bias generated by reads carrying reference vs. non-reference alleles. We designate the allele with the greater number of reads the strong allele and the other the weak allele. MARIO uses the ratio of the read counts between the two alleles to calculate an allelic reproducibility score (ARS) score, which factors in experimental replicates when available. ARS scores greater than 0.4 are used as a cutoff for selecting significant MARIO results, following the previous study [16].

## Declarations

### Ethics approval and consent to participate

Not applicable.

### Consent for publication

All authors read and approved the manuscript for review.

### Availability of data and materials

The NOMe-HiC datasets generated during the current study are available in the Gene Expression Omnibus (GEO) under the accession number GSE189158. Previously published data used for the analysis in this study are listed in Additional file 1: Table S1. The source code is publicly available with an MIT license at Zenodo (https://doi.org/10.5281/zenodo.7685935)[42] <u>and at BitBucket:</u> (https://bitbucket.org/dnaase/bisulfitehic/src/master/)[43].

### Authors’ contributions

Y.L. and L.W. conceived the study. H.F. and L.W. performed the experiment with input from Y.L. and L.J.M.. Y.L., H.Z., X.C. and M.T.W. performed computational analysis. Y.L., H.F., L.W., L.J.M., X.C. and M.T.W wrote the manuscript together. All authors read and approved the final manuscript.

## Acknowledgments

The authors thank Drs. Guoqiang Li and Bing Ren from the Ludwig Institute for Cancer Research and the University of California at San Diego, Dr. Fides Lay from Amgen Inc., and Dr. Raphael Kopan from Cincinnati Children’s Hospital Medical Center for helpful input. The authors also acknowledge the computational support from the Biomedical Informatics (BMI) high-performance computing cluster in CCHMC.

---

**Additional file 1** Supplementary materials.pdf: Supplementary Methods and Figures. Supplementary Methods and all Supplementary Figures in the study. Format: .pdf

**Additional file 2** Supplementary tables.xlsx: All the supplementary tables in the study. Format: .xlsx

