## Additional file 1 Supplementary Methods and Figures for "NOMe-HiC: joint profiling of genetic variants, DNA methylation, chromatin accessibility, and 3D genome in the same DNA molecule"

**Crosslinking, nuclei isolation, and DpnII digestion.** Cells were resuspended in ice-cold PBS at a concentration of 1M/mL and crosslinked with 1% formaldehyde (Thermo Scientific 28906) for 10min with gentle rotation at room temperature, followed by adding 0.2M Glycine and gently rotating in room temperature for 5min to stop crosslink. After washing, cells were suspended in Hi-C nuclei isolation buffer (10mM Tris-HCl Ph8.0, 10mM NaCl, 0.2% NP-40, 1x cOmplete protease inhibitor(Roche 11873580001)) supplemented with 1mM DTT, 0.02U/μL SUPERase• In™ RNase Inhibitor (Thermo AM2694) and 0.4U/μL RNaseOUT Recombinant Ribonuclease Inhibitor(Thermo Scientific 10777019) and incubate on ice for 1 hr. Nuclei were spun down at 2500g for 5 min at 4 ℃ and washed once with the above nuclei isolation buffer and spun down at 2500g for 5 min at 4 ℃. Then the nuclei membrane was permineralized with 50μL 0.5%SDS at 62 ℃ for 10 min. Permineralization was stopped by adding 25μL 10% Triton-X100 and 145μL $H_2O$. To digest the chromatin, 26.5μL NEB buffer3.1, 100U DpnII (NEB R0543M), 0.08U/μLSUPERase• In™ RNase Inhibitor, 0.16U/μL RNaseOUT Recombinant Ribonuclease Inhibitor, and 1mM DTT were added to the nuclei suspension and shaking overnight at 37 ℃.

**Biotin fill-in and proximal ligation.** DpnII was inactivated by incubation at 65 ℃ for 10 min and cool to room temperature for at least 10 min. Sticky ends were filled in with 0.05mM Biotin-1,4-dATP, 0.05mM dGTP, 0.05mM dCTP and 0.05mM dTTP by 0.13U/μL Klenow(NEB M0210L) at 37 ℃ for 90 min with 500 rpm shaking, supplemented with 0.02U/μLSUPERase• In™ RNase Inhibitor and 0.04U/μL RNaseOUT Recombinant Ribonuclease Inhibitor and 1mM DTT to prevent RNA degradation. Then proximal blunt-ends were ligated with 1.67U/μL T4 DNA Ligase (NEB M0202L) supplemented with 1X T4 DNA Ligase buffer, 1% Triton-X100 and 0.1mg/mL BSA (NEB B9000s) for 4 hr at room temperature with gentle shaking (300 rpm).

**M.cviPI treatment.** After ligation, nuclei were pelleted at 2500g for 5min at 4 ℃, followed by resuspension in 282μL 1x GpC buffer. Then nuclei suspension was treated with 50uL 4U/μL M.cviPI enzyme(NEB M0227L) supplemented with 1.5μL 32mM SAM, 150uL 1 M sucrose and 17μL 10xGpC buffer for 7.5min at 37 ℃. To enhance the efficiency of GpC methyltransferase footprint profiling, nuclei suspension was again treated with 25uL 4U/μL M.cviPI supplemented with 1.5μL 32mM SAM for 7.5min at 37 ℃. To stop the reaction, 973.5μL ice-cold PBS supplemented with 0.02U/μL SUPERase• In RNase Inhibitor and 0.04U/μL RNaseOUT Recombinant Ribonuclease Inhibitor and 1mM DTT were added. Then the nuclei suspension was split into two parts: 300μL for RNA-seq library preparation, 1200μL for whole-genome bisulfite library preparation.

**Nuclei RNA isolation.** Nuclei RNA was isolated with MagMAX FFPE DNA/RNA Ultra Kit (Thermo Scientific A31881) according to the protocol with minor modification. Briefly, 300μL Protease Digestion Buffer and 10uL Protease was added to 300μL nuclei suspension, and incubated at 55 ℃ overnight and 90 ℃ one hour to reverse crosslink the RNA. Then the suspension was cooled down to room temperature for 15 min. Nuclei RNA was captured by adding 20μL Dynabeads™ MyOne™ Silane (Thermo Scientific, 37002D) supplemented with 1000μL binding solution and 1250μL isopropanol, and shaking at 1000 rpm for 10 min at RT. The beads were collected by placing the sample-containing tube on a magnet for 5 min or until the supernatant was clear. Then the beads were washed once with a 500uL RNA wash buffer and once with a 500μL wash solution. DNA was digested by resuspending the Saline beads in 20μL DNase, 10μL DNase buffer, and 70μL $H_2O$ and incubating at RT for 20 min. To recapture the RNA, 200μL binding buffer and 250μL isopropanol were added and incubated at RT for 10 min with 1000 rpm shaking. The beads were again washed once with the RNA wash buffer and twice with the wash solution. After briefly air-drying the beads, nuclei RNA was eluted with 30μL nuclease-free $H_2O$.

**RNA-seq library preparation.** 10ng nuclei RNA was used for library preparation using SMARTer® Stranded Total RNA-Seq Kit v2 - Pico Input Mammalian - 96 Rxns (TAKARA 634413) kit.

**Reverse crosslink, DNA precipitation, and sonication.** Nuclei DNA was first reverse crosslinked by adding 50μL 20 mg/mL proteinase K and 120μL 10% SDS, then incubating at 55 ℃ for 30 min. The transcription factor was disload by adding 130μL of 5M NaCl and then incubated at 68 ℃ for 4 hrs. Tubes were cooled to room temperature, and nuclei DNA was precipitated at -20 ℃ overnight with 1.6x EtOH and 0.1x NaAc. The next morning, nuclei DNA was collected by spin down at max speed for 15 min at 4 ℃, and washed twice with fresh 80% EtOH, then dissolved in 130μL 10mM Tris-HCl ph8.0. The DNA was sonicated to an average of 400bp with Covaris M220 using the following parameters: peak power 30, duty factor 10, cycles/burst 200, duration 60s. A 1x AMPure size selection was performed to get rid of the small fragments, and nuclei DNA was then suspended in 300μL 10mM Tris-HCl ph8.0.

**Biotin pull-down, end-repair, dA-tailing, and library preparation.** Dynabeads My One T1 Streptavidin beads (Invitrogen 65602) was washed once with 1x tween wash buffer (5mM Tris-HCl ph7.5, 0.5mM EDTA, 1M NaCl, 0.05% Tween-20) and suspended in 300μL 2x binding buffer (10mM Tris-HCl ph7.5, 1mM EDTA and 2M NaCl). Then biotin pull-down was performed by rotation at room temperature for 15min. After that, beads-DNA was washed twice with a 1x tween wash buffer at 55 ℃ with mixing. Then end-repair was performed, and unligated fragments were removed with 0.5mM dNTP (Thermo R1122), 0.5U/μL T4 PNK (NEB M0201L), 0.12U/μL T4 DNA polymerase (NEB M0203L), 0.05U/μL Klenow(NEB M0210L) in 100μL 1x T4 DNA ligase buffer at room temperature for 30 min with 300 rpm mixing. DNA was washed twice at 55 °C with mixing. dA-tailing was performed 0.5mM dATP, 0.25U/uL Klenow(exo-) (NEB M0212L) in 100μL 1xNEB buffer 2 at 37 ℃ for 30 min.  DNA was washed twice at 55 °C with mixing. 0.9μM truncated cytosine-methylated X-gene Universal Stubby Adapter (IDT) was added by Quick Ligase in 50μL 1x Quick Ligase buffer at RT for 15min. Nuclei DNA was washed twice with 1x tween wash buffer at 55 ℃ followed by washing twice with 100μL 10mM Tris-HCl pH8.0. Finally, DNA binding streptavidin beads were resuspended in 20μL 10mM Tris-HCl phH8.0.

**Bisulfite conversion.** Spike in 0.1% 400bp stubby ligated unmethylated lambda DNA into each DNA sample before bisulfite conversion. Bisulfite conversion was performed with EZ DNA Methylation-Gold Kit (Zymo D5006). DNA was separated from streptavidin beads after CT conversion.

**Library amplification and sequencing.** The Bisulfite converted library was amplified using 100ng input with KAPA HiFi Uracil+ ReadyMix(KAPA KK2801), 0.5μM truncated TruSeq™–Compatible Indexing Primer, 2.5mM $MgCl_2$. The following cycles were performed: 98 ℃ 45 sec; 98 ℃ 15 sec, 60 ℃ 30 sec, 72 ℃ 30 sec (10~15 cycles); 72 ℃, 1 min. After pooling and quality control, sequencing was performed at HiSeq-XTen or NovaSeq S6000 platform with 150 paired ends.

### Supplementary Figures

**Supplementary Figure 1**

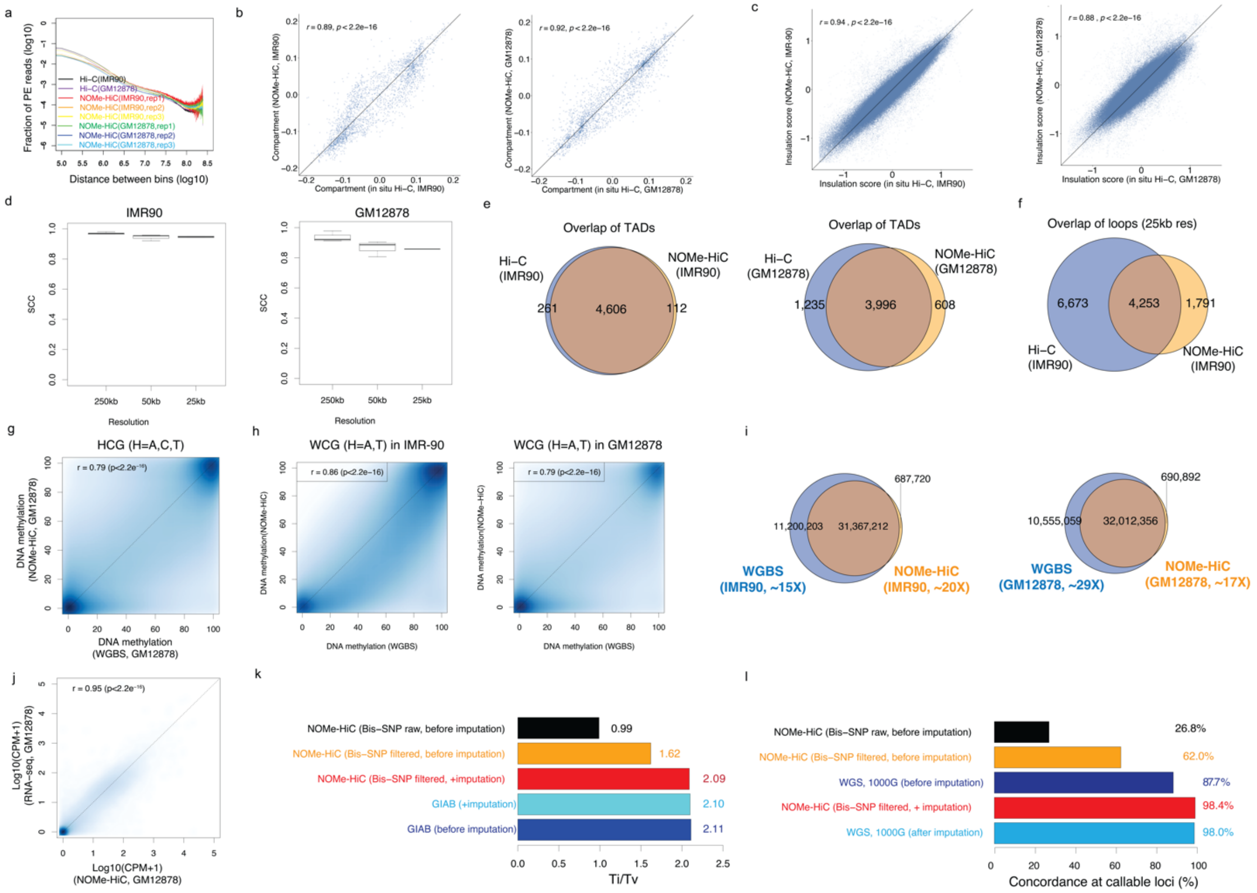


**Fig. S1. NOMe-HiC generates high-quality multi-omics data at different cell types**. (a). Comparison of contact frequency distance decay curve obtained from *in situ* Hi-C and NOMe-HiC data at IMR-90 and GM12878. (b). Comparison of compartment score (500kb resolution) obtained from *in situ* Hi-C and NOMe-HiC data at IMR-90 (left) and GM12878 (right). (c). Comparison of insulation score (10kb bin) obtained from *in situ* Hi-C and NOMe-HiC data at IMR-90 (left) and GM12878 (right). (d). Matrix similarity between *in situ* Hi-C and NOMe-HiC measured by stratum adjusted correlation coefficient (SCC) from HiCRep at different resolutions in IMR-90 (left) and GM12878 (right). (e). Overlap of TADs obtained from *in situ* Hi-C and NOMe-HiC data at IMR-90 (left) and GM12878 (right) by TopDom (50kb resolution). (f). Overlap of chromatin loops (25kb resolution) obtained from *in situ* Hi-C and NOMe-HiC data at IMR-90 by MUSTACHE. (g). The methylation concordance with WGBS at HCG (H=A, C, or T) sites in GM12878. (h). The methylation concordance with WGBS at WCG (H=A, or T) sites in IMR-90 (left) and GM12878 (right). (i). The number of HCG sites covered by *in situ* Hi-C and NOMe-HiC data at IMR-90 (left) and GM12878 (right). (j). Scatterplot of the concordance at the gene level with total RNA-seq (ENCODE) in GM12878. (k). Ti/Tv ratio at gold standard data (GIAB) before or after imputation and at Bis-SNP results (filtered or raw) before and after imputation from NOMe-HiC in NA12878. (l). The percentage of concordant SNPs with the ground truth (GIAB) at the callable sites (with reads covered and genotypes called) in WGS and NOMe-HiC (by Bis-SNP, filtered or raw) in NA12878 before or after the imputation.

**Supplementary Figure 2**

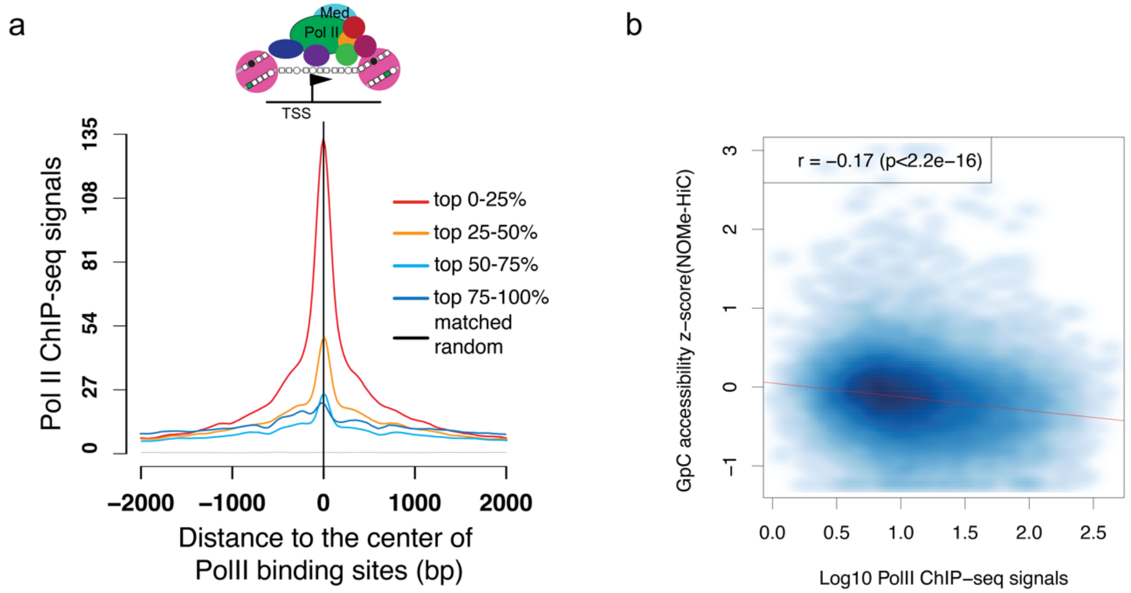


**Fig. S2. GpC methyltransferase footprint level is correlated with PolII binding signals.** (a). The PolII ChIP-seq signal strength at different quantiles near promoters. (b) The scatterplot of -log10pvalue of PolII ChIP-seq signals and z-score transformed GCH methylation level at PolII ChIP-seq IDR peaks obtained in IMR-90 cell lines from ENCODE.

**Supplementary Figure 3**

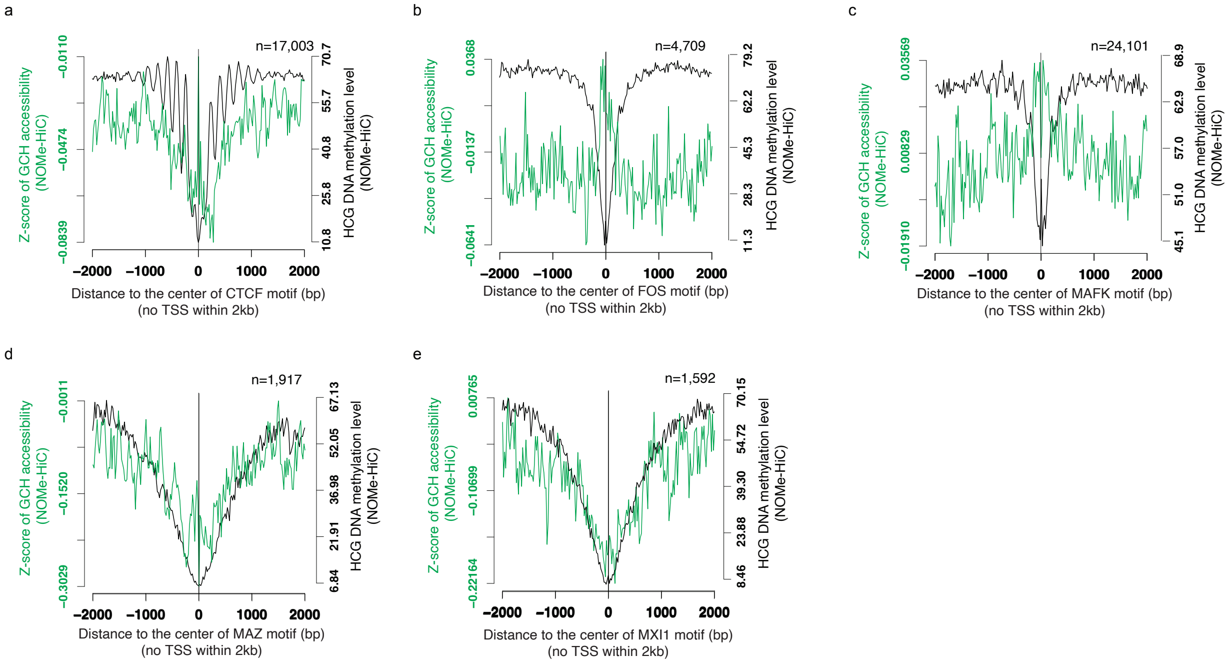


**Fig. S3. Average GCH methyltransferase footprint and HCG methylation level around distal (a). CTCF (b). FOS (c). MAFK (d). MAZ (e). MIX1 binding sites in IMR-90**. The TF binding sites are characterized by TF ChIP-seq in IMR-90.

#### Supplementary Figure 4

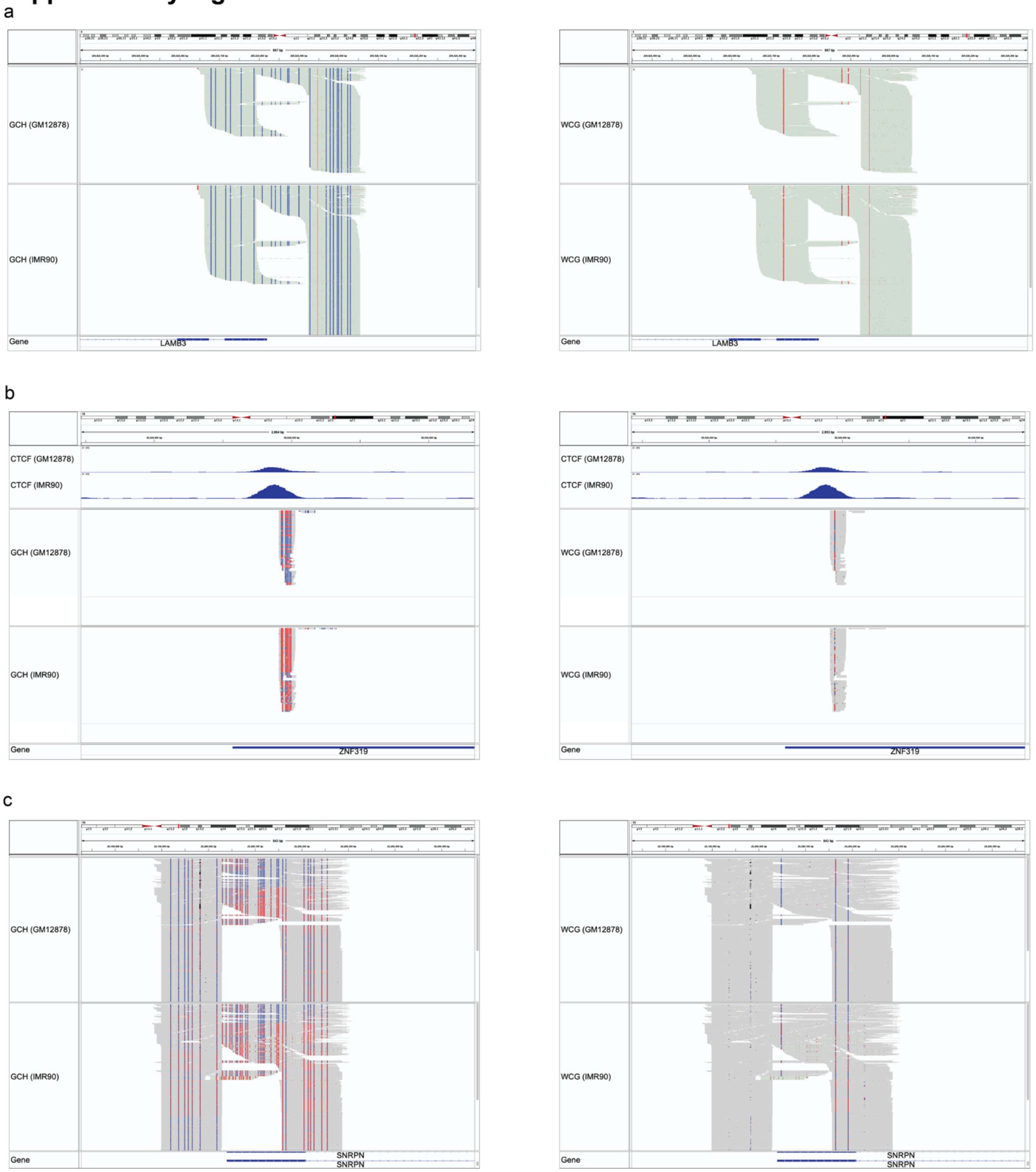


**Fig. S4. GCH (left) and WCG (right) methylation level measured by locus-specific amplification at (a). LAMB3, (b) CTCF peak (c).SNRPN site of NOMe-HiC libraries from IMR-90 and GM12878 cell lines.**

#### Supplementary Figure 5

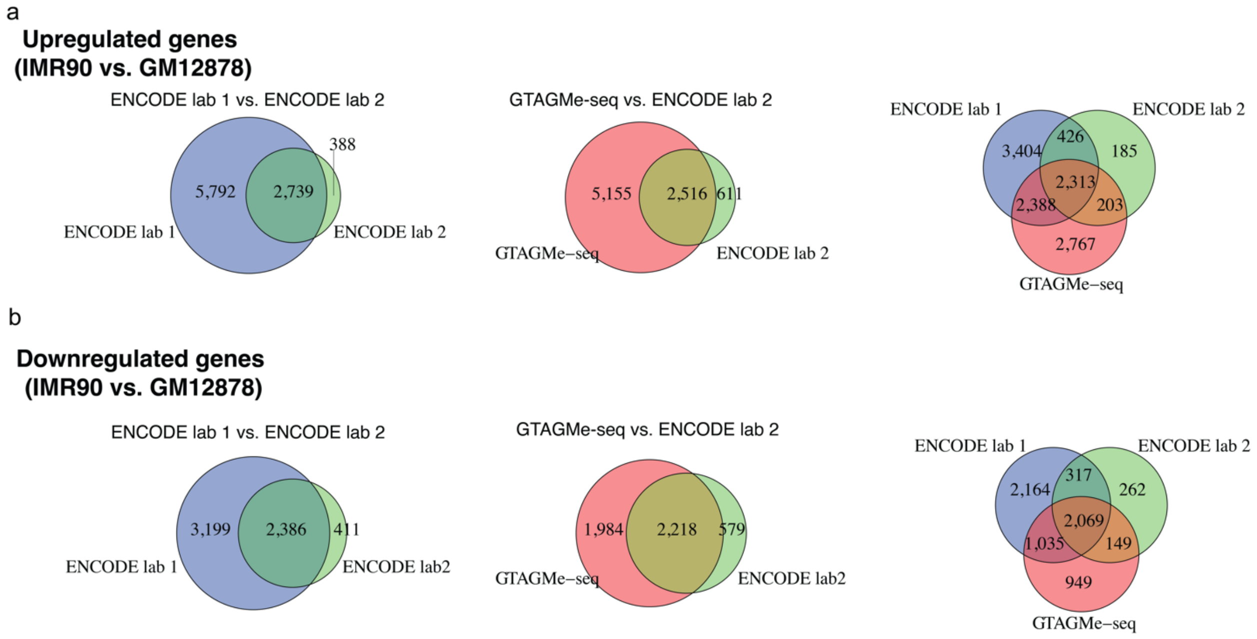


**Fig. S5. NOMe-HiC reveals similar sets of differential expressed genes as total RNA-seq**. (a). The overlap of significantly upregulated genes between IMR-90 and GM12878 that are characterized by total RNA-seq from two different labs in ENCODE (left), or characterized by NOMe-HiC and total RNA-seq from ENCODE (middle), or characterized by three datasets (right). (b). The overlap of significantly downregulated genes between IMR-90 and GM12878 that are characterized by total RNA-seq from two different labs in ENCODE (left), or characterized by NOMe-HiC and total RNA-seq from ENCODE (middle), or characterized by three datasets (right).

**Supplementary Figure 6**

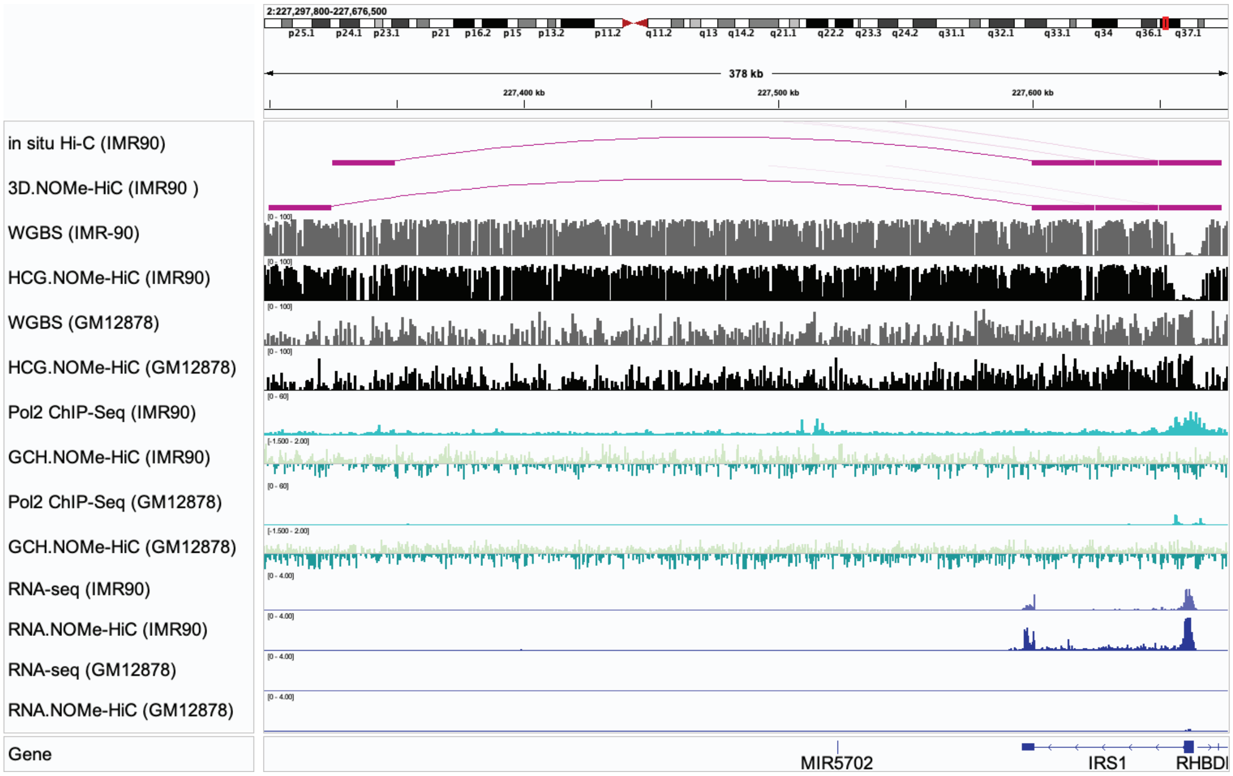


**Fig. S6. Example plot of NOMe-HiC**.

### Supplementary Figure 7

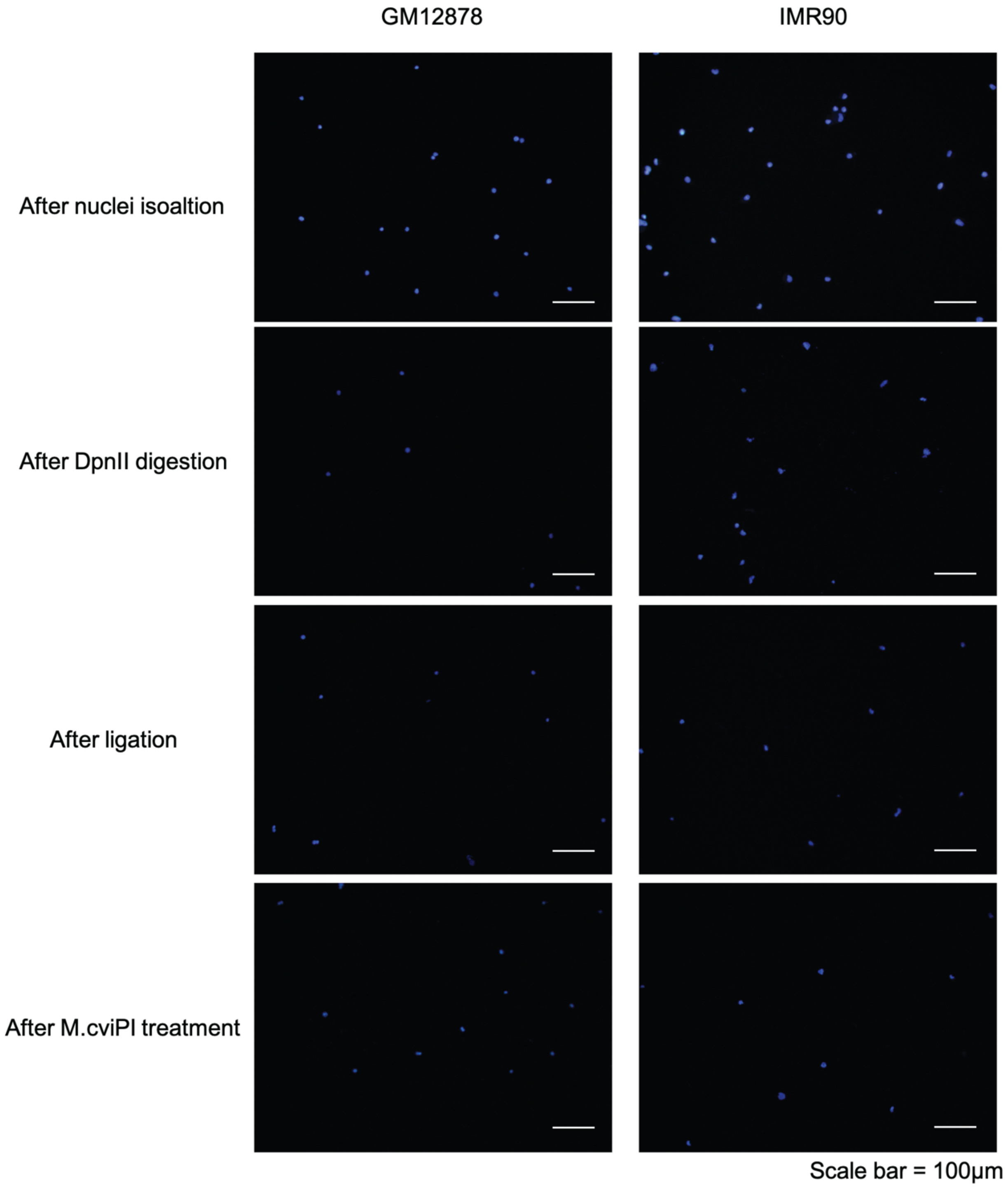


**Fig. S7. Nuclei status at each step of NOMe-HiC library preparation**.

**Supplementary Figure 8**

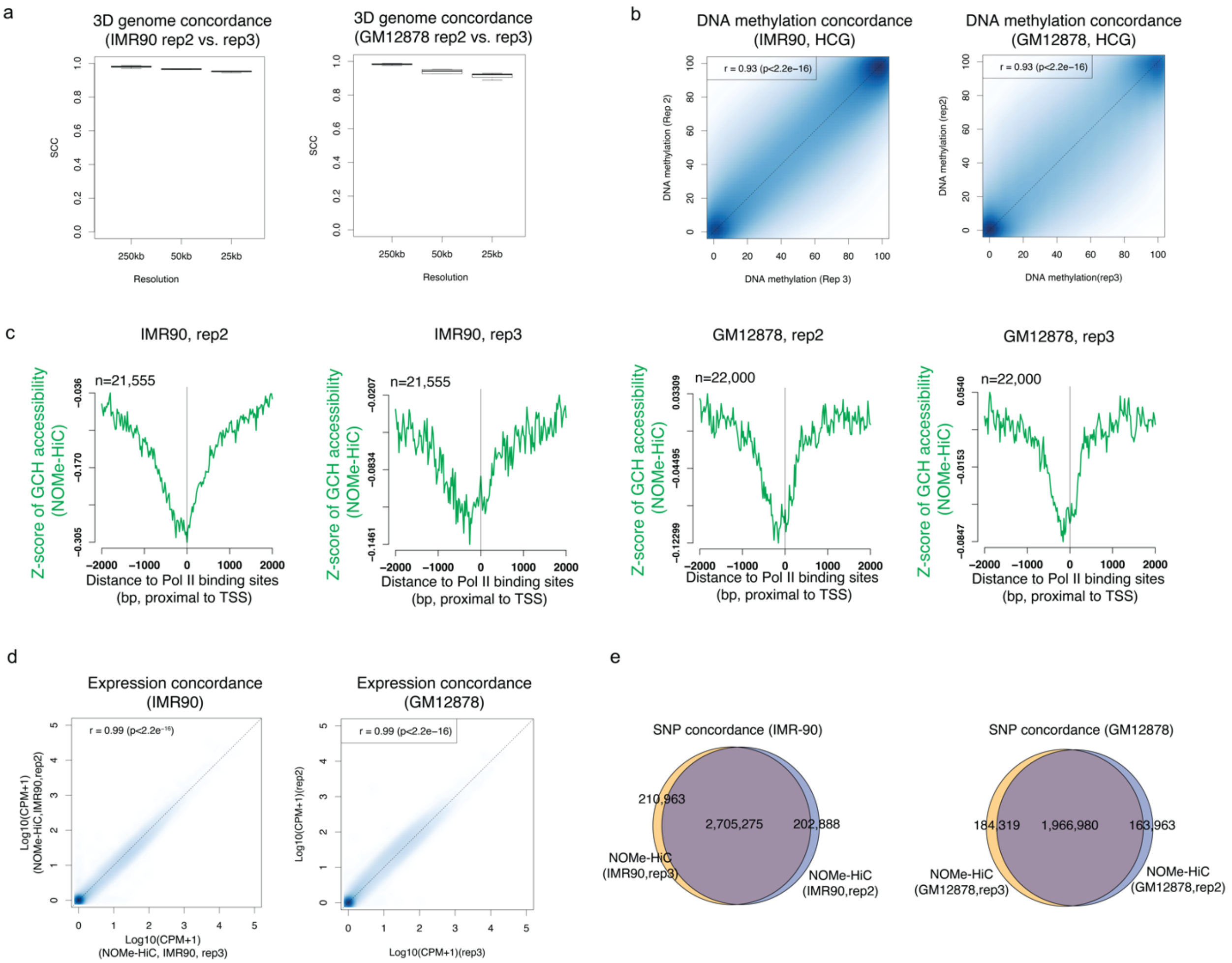


**Fig. S8. NOMe-HiC generated highly reproducible multi-omics data across biological replicates at two cell lines**. (a). 3D genome similarity between two biological replicates in NOMe-HiC measured by stratum adjusted correlation coefficient (SCC) from HiCRep at different resolutions in IMR-90 (left) and GM12878 (right). (b). The methylation concordance between two biological replicates in NOMe-HiC at HCG sites in IMR-90 (left) and GM12878 (right). (c). Z-score transformed GCH accessibility near PolII binding sites in two biological replicates in NOMe-HiC at HCG sites in IMR-90 (left) and GM12878 (right). (d). Scatterplot of the concordance at the gene level between biological replicates in NOMe-HiC data at IMR-90 (left) and GM12878 (right). (e). SNP concordance between biological replicates in NOMe-HiC data at IMR-90 (left) and GM12878 (right).

### Supplementary Figure 9

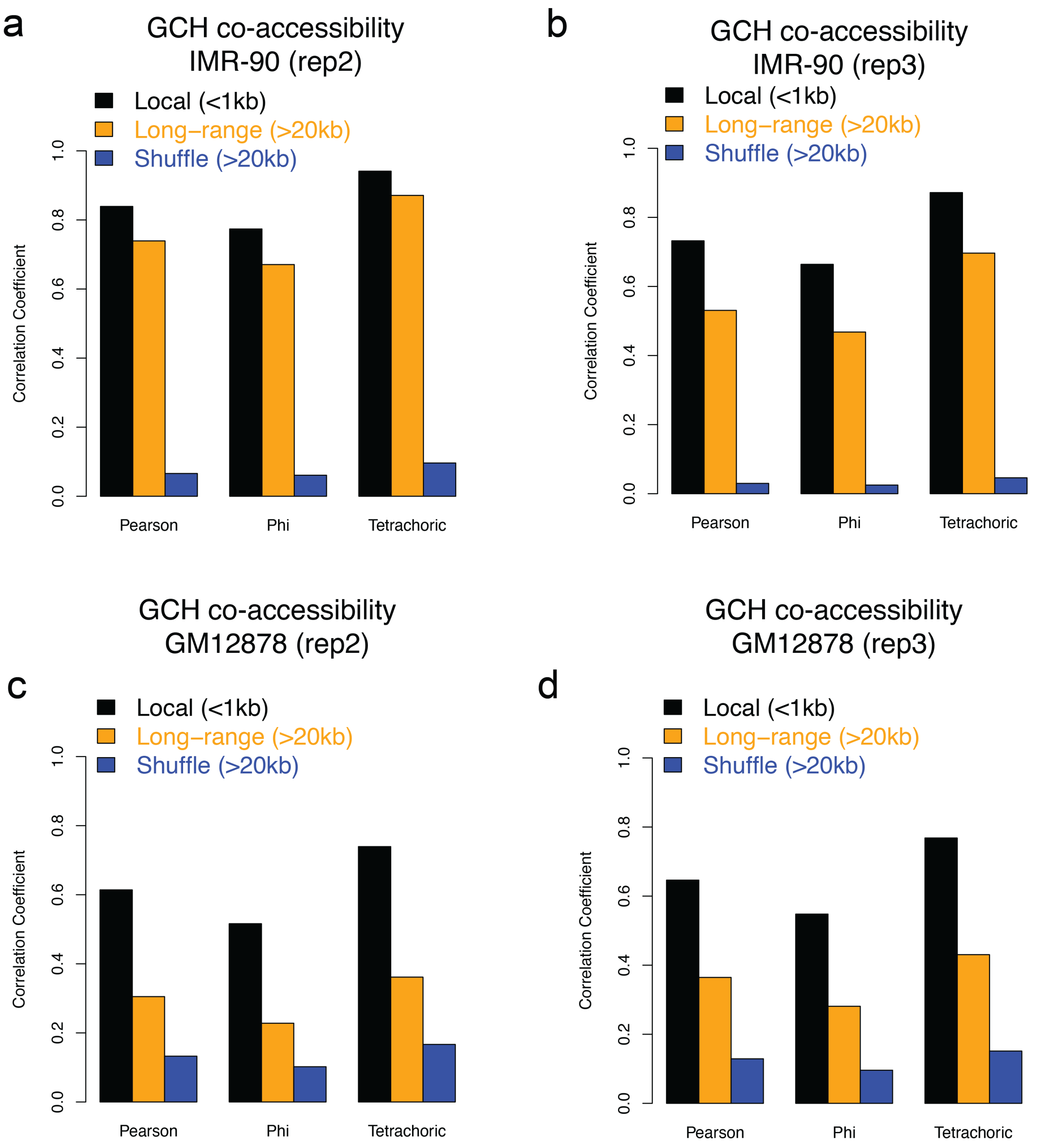


**Fig. S9. NOMe-HiC generated reproducible long-range GCH accessibility concordance across biological replicates at two cell lines**. Average GCH methyltransferase concordance level, measured by Pearson Correlation Coefficient, Phi Correlation Coefficient, and Tetrachoric correlation, across all the chromatin loops anchor regions (>20kb distance between anchors, orange color), their matched local regions (utilize the read pairs from the same anchor and within 1kb distance, black color), and matched shuffled read pairs from the same chromatin loops anchor regions (>20kb distance between anchors, blue color) at (a). IMR-90 rep2 (n=377,640 for short-range pairs, n=46,576 for long-range pairs), (b). IMR-90 rep3(n=378,359 for short-range pairs, n=78,008 for long-range pairs), (c). GM12878 rep2(n=703,529 for short-range pairs, n=10,650 for long-range pairs), and (d). GM12878 rep3(n=693,107 for short-range pairs, n=15,868 for long-range pairs).

### Supplementary Figure 10

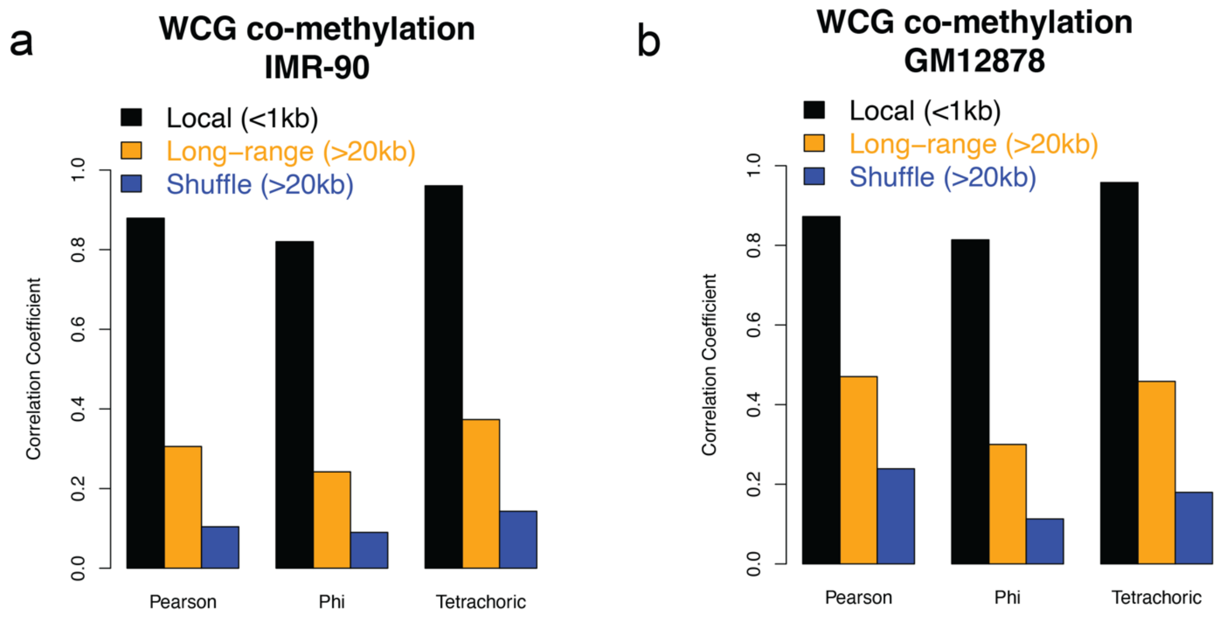


**Fig. S10. NOMe-HiC reproduced the long-range methylation concordance at two cell lines**. Average WCG methylation concordance level, measured by Pearson Correlation Coefficient, Phi Correlation Coefficient, and Tetrachoric correlation, across all the chromatin loops anchor regions (>20kb distance between anchors, orange color), their matched local regions (utilize the read pairs from the same anchor and within 1kb distance, black color), and matched shuffled read pairs from the same chromatin loops anchor regions (>20kb distance between anchors, blue color) at (a). IMR-90. and (b). GM12878.

#### Supplementary Figure 11

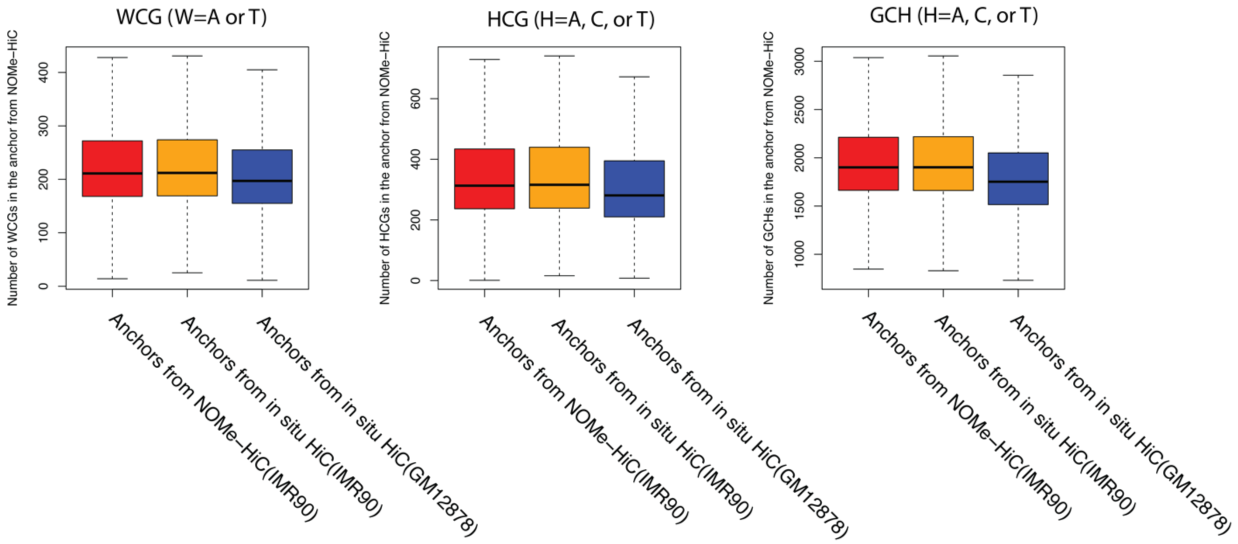


**Fig. S11. The boxplot of the number of WCG, HCG, and GCH at loop anchors in NOMe-HiC.**

### Supplementary Figure 12

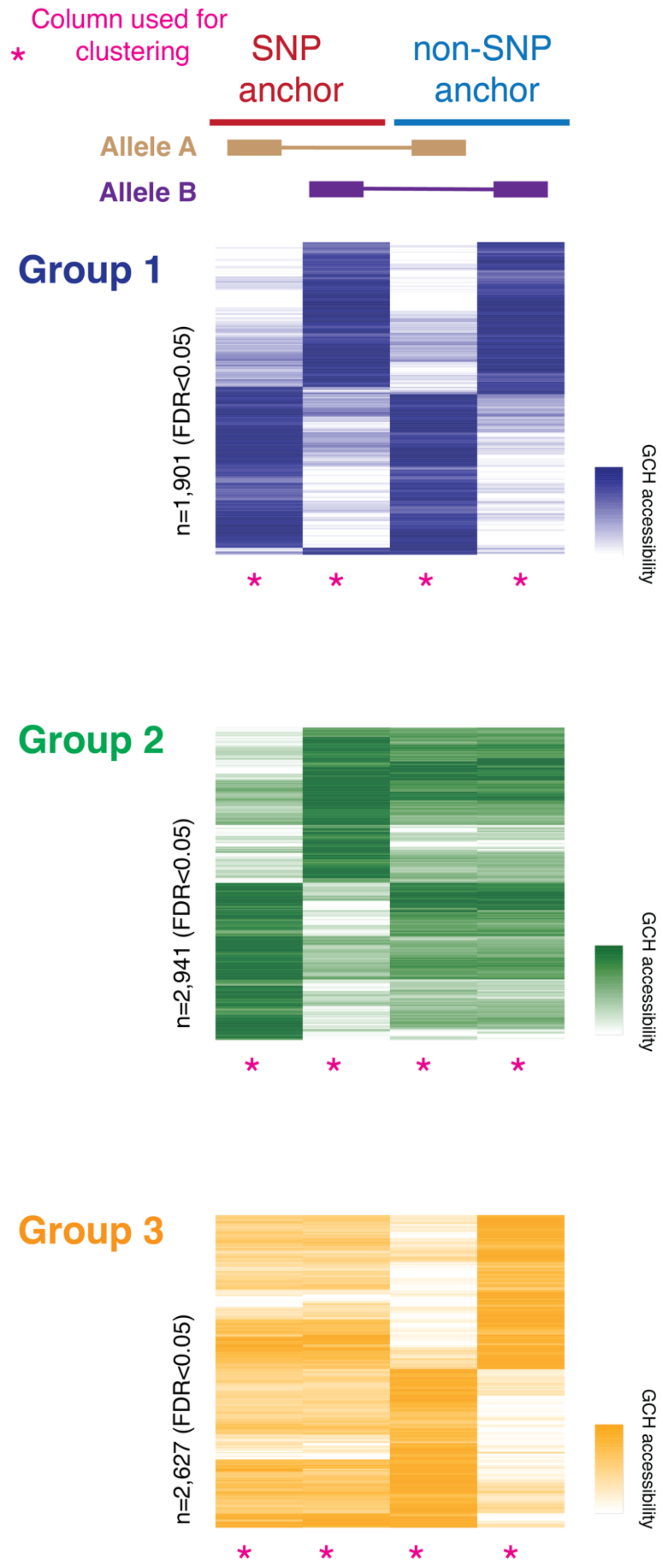

**Fig. S12. NOMe-HiC reveals the long-range allele-specific GCH methyltransferase footprint**. The rows of the heatmap are clustered by all four columns (marked with magenta star *) instead of one column in the main Figure 4.

#### Supplementary Figure 13

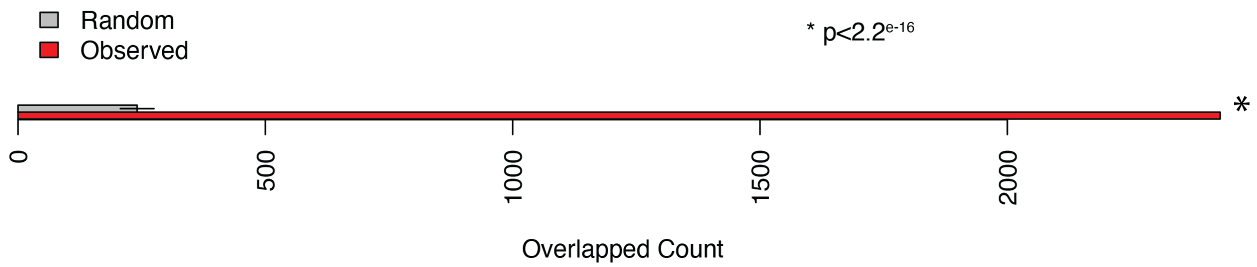


**Fig. S13. The enrichment of allele-specific TF binding sites at the SNP anchors of allele-specific GCH methyltransferase footprint loci.** P value is calculated by the Fisher exact test with the random permutated overlap numbers.

**Table S1:** The list of public data used in the study.

**Table S2:** Sequencing summary statistics in NOMe-HiC (DNA).

**Table S3:** Sequencing summary statistics in NOMe-HiC (RNA).

**Table S4:** Three groups of long-range allele specific GpC methyltransferase footprint regions.

**Table S5:** LD blocks overlapped with the SNP anchors in group 3 Allele-specific GpC methyltransferase footprint.
